# On Using Local Ancestry to Characterize the Genetic Architecture of Human Phenotypes: Genetic Regulation of Gene Expression in Multiethnic or Admixed Populations as a Model

**DOI:** 10.1101/483107

**Authors:** Yizhen Zhong, Minoli Perera, Eric R. Gamazon

**Affiliations:** Department of Pharmacology, Northwestern University Feinberg School of Medicine, Chicago, IL, USA; Vanderbilt Genetics Institute and Division of Genetic Medicine, Vanderbilt University School of Medicine, Nashville, TN, USA; Clare Hall, University of Cambridge, Cambridge, UK

**Keywords:** admixture, eQTL, local ancestry, transcriptome, mixed models, heritability, omics, population structure

## Abstract

**Background:** Understanding the nature of the genetic regulation of gene expression promises to advance our understanding of the genetic basis of disease. However, the methodological impact of use of local ancestry on high-dimensional omics analyses, including most prominently expression quantitative trait loci (eQTL) mapping and trait heritability estimation, in admixed populations remains critically underexplored.

**Results:** Here we develop a statistical framework that characterizes the relationships among the determinants of the genetic architecture of an important class of molecular traits. We estimate the trait variance explained by ancestry using local admixture relatedness between individuals. Using National Institute of General Medical Sciences (NIGMS) and Genotype-Tissue Expression (GTEx) datasets, we show that use of local ancestry can substantially improve eQTL mapping and heritability estimation and characterize the sparse versus polygenic component of gene expression in admixed and multiethnic populations respectively. Using simulations of diverse genetic architectures to estimate trait heritability and the level of confounding, we show improved accuracy given individual-level data and evaluate a summary statistics based approach. Furthermore, we provide a computationally efficient approach to local ancestry analysis in eQTL mapping while increasing control of type I and type II error over traditional approaches.

**Conclusion:** Our study has important methodological implications on genetic analysis of omics traits across a range of genomic contexts, from a single variant to a prioritized region to the entire genome. Our findings highlight the importance of using local ancestry to better characterize the heritability of complex traits and to more accurately map genetic associations.

## Background

Greater understanding of the genetic determinants of high-dimensional molecular traits, such as the prominent example of eQTL, promises to advance our understanding of the genetic architecture of complex traits [1, 2]. Since the majority of trait-associated variants identified by genome-wide association studies (GWAS) reside in non-coding regions [3], eQTL data provide an important resource to elucidate the underlying mechanisms of these non-coding variants by linking them to gene expression [1]. In addition, heritability estimation, i.e., determining the trait variance explained by regulatory variants, may provide important insights into the genetic architecture of gene expression traits. However, to date, eQTL mapping and heritability analysis have been conducted primarily in populations of European ancestry, and omics data in recently admixed populations, such as African Americans (AAs), that are disproportionately affected by a variety of complex diseases, are lacking [4-7], limiting our understanding of the genetic basis of trait variance in human populations. Populations of African descent have greater genetic variation and less extensive linkage disequilibrium (LD), which may restrict the generalizability of genetic associations identified in non-African populations to AAs [8, 9]. Importantly, the impact of the admixed genome structure on eQTL mapping and heritability estimation has not been adequately studied.

The eQTL (regulatory) effect on gene expression is typically modeled (using linear regression) assuming additive effect of genetic variation on gene expression [10]. The resulting association analysis tests only the correlation between genotype and phenotype instead of testing for causal effects and is easily subject to confounding from population structure. The chromosomes of AAs comprise of mosaic regions of different ancestral origins, resulting in two types of population structure that may be present in genetic association analyses [11]. One arises from *global ancestry*, which reflects the admixture proportions of the (previously isolated) ancestral populations (primarily African and European, though a relatively small proportion of Native American ancestry [12] may also be present) and is typically estimated with the first principal component (PC) derived from genome-wide genotype data, which separates the European and African ancestral populations [13]. (The assumption of a small number of ancestral populations is often made for methodological and computational convenience.) The PCs have been shown to have a geographic interpretation and their use widely adopted due to computational efficiency [14, 15]. Mixed models incorporate the pairwise genetic similarity between every pair of individuals in the association mapping and have been effectively deployed to correct for population structure, family structure, and cryptic relatedness [16, 17], but until recently, mixed model approaches have been too computationally intensive for eQTL mapping.

Population structure in association studies of an admixed population may also arise from *local ancestry*, which is the number of inherited alleles (0, 1 or 2) from each ancestral population at a particular locus [18]. Local ancestry may vary across the genome as well as across individuals, even of similar global ancestry [13], at any given locus. As a large proportion of gene expression phenotypes have been found to be differentially expressed between Africans and Europeans [4], increased spurious eQTL associations (false positives) could arise, leading to pseudo-associations that are not driven by the genetic variants being tested but, instead, by their local ancestral backgrounds. Studies that explore the methodological importance of local ancestry in genetic association analyses have been limited to a small number of highly polygenic traits [19, 20]. Incorporating local ancestry into eQTL mapping, which tests associations between millions of SNPs and thousands of genes, has been too computationally intensive.

Heritability estimation is usually performed using linear mixed models (LMM) but has been conducted primarily in ancestrally homogeneous populations. LD Score Regression (LDSR) [21] is a summary statistics based approach to estimating heritability and confounding, but its applicability to studies involving admixed individuals has not been investigated. Heritability of gene expression traits ^h^as been characterized by a more sparse genetic architecture [22] and by an *a priori* functionally relevant (cis) region in contrast to polygenic complex traits, suggesting a greater role for local ancestry than global ancestry. Local ancestry may be determined by a range of factors, including population demographic history (e.g., migration, population bottleneck, etc.), which can shape complex admixture dynamics (e.g., as trans-atlantic migration has impacted the local ancestry of African Americans). The impact of the use of local ancestry on estimating the heritability of gene expression traits is thus a critical gap in our understanding of their genetic architecture. Furthermore, high-dimensional omics studies provide an opportunity to assess, more comprehensively, the contribution of local ancestry to human phenotypic variation through joint analysis of thousands of molecular traits.

Here we provide a statistical framework to analyze the relationships among the proportion of variance explained (PVE) by genetic variation (PVE_*g*_), PVE by local ancestry (PVE_*l*_), global ancestry, and degree of population differentiation at causal regulatory variants for gene expression traits in admixed populations. We performed a comprehensive analysis of the variation in gene expression explained by local ancestry versus global ancestry. We analyzed the impact of the use of local ancestry on eQTL mapping and heritability estimation through extensive simulations and the application of our approach in a transcriptome dataset in an admixed population as well as data from the (GTEx) project [2] consisting of multiethnic individuals. We develop an efficient approach to eQTL mapping in an admixed population, demonstrating that use of local ancestry can substantially improve mapping of genetic associations. We demonstrate that our approach shows improved control of type I error rate as well as increased statistical power compared with a global ancestry adjustment approach in eQTL mapping, and find greater replication rate for eQTLs specific to our approach. Finally, we propose a novel method for heritability estimation in admixed populations, opening new avenues for research into the genetic architecture of complex traits.

## Results

### Relationship among effect of genetic variation on gene expression, variance explained by local ancestry, population differentiation, and global ancestry

Gene expression may differ in its genetic architecture from a complex disease or general quantitative trait in several crucial ways, including the importance of the local (cis) region and the potential for a large sparse genetic component [23]. In the case of an admixed population, we hypothesize that the ancestry background near the gene of interest may have a primary importance, with local ancestry potentially explaining a greater proportion of transcriptional variation than global ancestry. We therefore consider these key features in modeling the trait variance explained by local ancestry and genetic variation (see **Methods**).

First, we assume the simplest case of a single causal variant, as sometimes assumed in certain eQTL analyses (such as fine mapping and single-variant association tests). We define the population genetic parameter, *F*_*st,f*_ at the variant *f* in terms of the allele frequencies *p*_*f*,1-_ and *p*_*f*,.0_ (in the ancestral populations 1 and 0) and the genotype variance 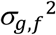 as follows:

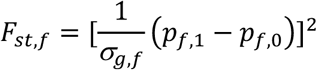

We then obtain the following (see **Methods** for derivation):

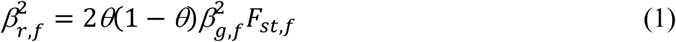

This expression relates, for a given causal eQTL, the effect explained by local ancestry 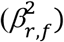 with the effect of the genetic variant on gene expression 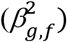, global ancestry (*θ*), and the degree of population differentiation (*F*_*st,f*_).

We extend equation (1) to the case of multiple casual eQTL variants in the cis region of a gene, as would be relevant for heritability estimation assuming allelic heterogeneity. We obtain the following inequality (see **Methods**):

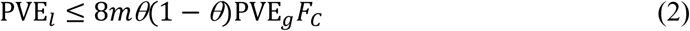

where 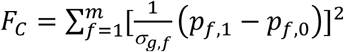 is the total extent of population differentiation at causal eQTL variants. This provides an upper bound on the trait variance explained by local ancestry (PVE_*l*_) in terms of the aggregate genetic effect (PVE_*g*_), the magnitude of population differentiation of the causal regulatory variants (*F*_*c*_), and the degree of polygenicity of the gene expression trait (captured by the number of causal eQTLs, *m*). We confirmed inequality (2) using simulations across a range of genetic architectures (see **Methods** and Additional file 1: **Table S1**).

### Implications of statistical model

From equation (1), potential sources of bias in the estimate of *β*_*g,f*_ include all the remaining parameters, which are local parameters and indeed depend on *f*. In addition to the level of admixture, uncertainty in local ancestry estimation, and the degree of population differentiation may contribute to bias. Importantly, global ancestry adjustment ignores local heterogeneity in LD pattern and ancestral population allele frequency difference across the genome. (We investigate the single-variant case extensively below when we evaluate the use local ancestry in mapping genetic associations.) From inequality (2), these consequences follow:

1. The parameters *θ* (a characteristic of the population) and *F*_*st,f*_ (a population genetic parameter that measures genetic distance between the ancestral populations) are *a priori* unrelated to the phenotype (gene expression) while PVE_*g*_ and the polygenicity parameter *m* are specific to the phenotype. The population differentiation statistic *F*_*stc*_ used by a recent study [24] assumes a highly specific genetic architecture and incorporates the trait-dependent weight 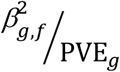 at the variant *f*, thus varies by phenotype for each variant, and is consequently not a purely population genetic parameter. For eQTL studies involving thousands of (gene expression) phenotypes with varying levels of polygenicity and potentially displaying a range of genetic architectures, we wanted to utilize a *phenotype-independent* measure of genetic distance between the ancestral populations at each variant, which determined our model and led to inequality (2). Of course, summing the genetic distance over all causal eQTL variants for the specific gene expression trait introduces phenotype dependence. Nevertheless, assuming shared eQTLs across populations (though with possibly different allele frequencies), this framework would facilitate more straightforward population comparisons by disentangling the contribution of the population genetic parameters from that of the phenotype-dependent variables.
2. Since the maximum of θ(1 - θ) is 0.25, the quantity 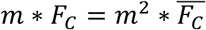 in inequality (2) determines whether local ancestry (in the cis region) explains less of the transcriptional variation than genetic variation. In particular, if *F*_*c*_ < 1/(2*m*), local ancestry would explain less of the variation in gene expression. Now the quantity 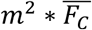 is linear in the mean level of differentiation at the gene but quadratic in the degree of polygenicity, indicating that characterization of a gene expression trait as sparse or polygenic has important implications for assessing the variation explained by local ancestry and by genetic variation.
3. If we assume (a) a highly polygenic architecture for a gene expression trait with each causal variant contributing only a modest proportion that depends only on the total number of contributing variants, i.e., 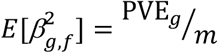, and (b) the independence of the causal variant contribution to trait variance and degree of population differentiation (i.e., independence of 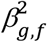 and *F*_*st,f*_), we obtain (see **Methods**):

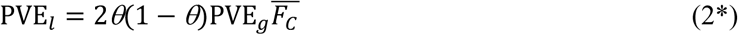

where 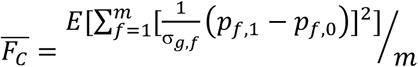. Equation (2*) therefore provides an estimate 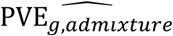 for PVE_*g*_, which can be viewed as the “expected heritability” in the presence of admixture, departure from which may yield additional insights into genetic architecture (see **Methods**). The condition 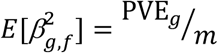 is an assumption about the causal eQTL effects being drawn from a single (Gaussian) distribution with the given expected value or mean. We note that the assumption of a single distribution of effect sizes may be a reasonable one for all cis effects, but trans effects may plausibly require a different distribution. Similarly, a single effect size distribution may not hold for both common and rare regulatory variants. The condition is thus a strong assumption about how heritability is distributed across the cis region of the gene, with its assignment of causal effects from the same distribution independently of LD. Under the two assumptions of polygenicity and independence, we get PVE_*l*_ ≤ 0.50(PVE_*g*_) from inequality (2), indicating that the variance explained by local ancestry would be less than that explained by local genetic variation. We emphasize that inequality (2) holds for a wide range of genetic architectures but equation (2*) assumes strict constraints on the genetic architecture.

We note that the statistical model applies more broadly to the analysis of trait variance explained by local ancestry and genetic variation in studies of the proteome, the methylome, and other types of omics data. Furthermore, inequality (2) and equation (2*) characterize the expected PVE_*g*_ in the presence of admixture, and violation of these relationships may well indicate the presence of stratification or, in the case of equation (2*), violation of at least one of the two assumptions of polygenicity and independence (see **Methods**).

### Local ancestry and mapping genetic associations

We sought to investigate the importance of use of local ancestry for eQTL mapping in an admixed population (Joint-GaLA-QTLM; see **Methods**). We plotted the empirical distribution of the maximum *β*_*l*_ of each gene and found, for a number of genes, that a large proportion of the variance in expression can be explained by the local ancestry at a single variant in *both* NIGMS and GTEx (NIGMS: 36 out of 4,595 genes, FDR<0.10, **Figure 1A**; GTEx whole blood: 3,129 out of 19,432 genes, FDR<0.10, **Figure 1B**), suggesting that the confounding due to local ancestry might exist not only in studies of recently admixed populations but also in studies with multiethnic samples.

**Figure 1.**
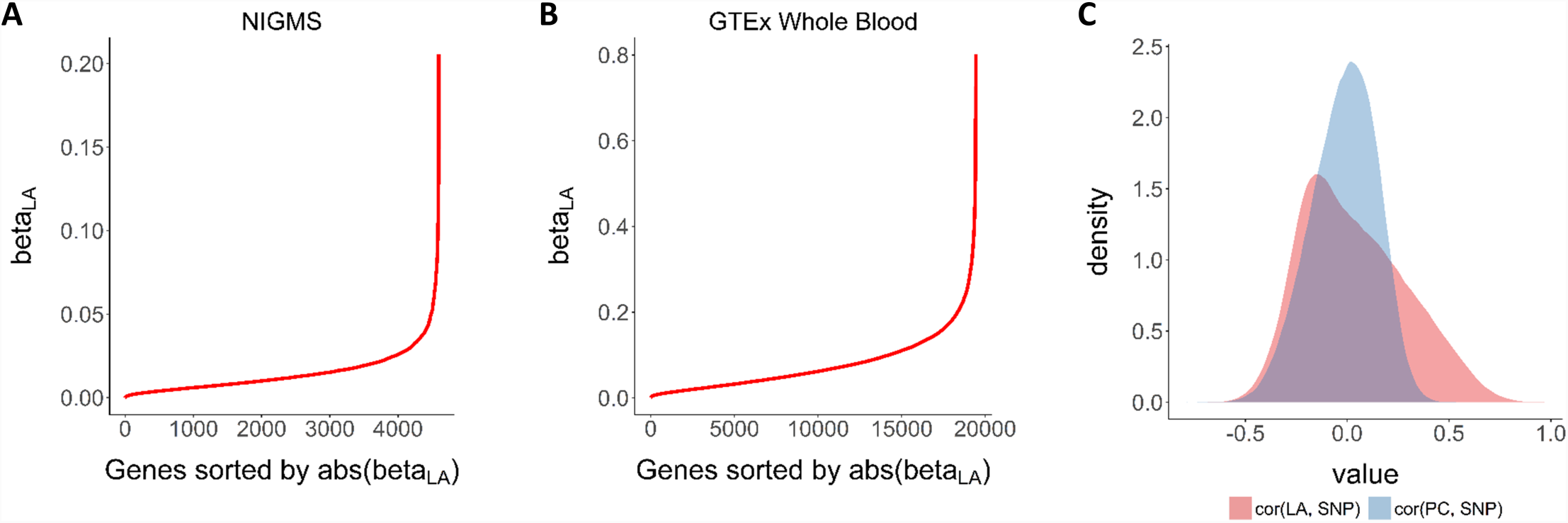
Effect explained by local ancestry and eQTL association test. We evaluated the effect explained by local ancestry and the correlation between local ancestry and genotype. **A, B.** Empirical distribution of the maximum effect size of local ancestry for each gene expression trait in NIGMS dataset (admixed samples, **1A**) and in the GTEx whole blood dataset (multiethnic samples, **1B**), showing a large effect for a substantial number of genes. **C.** Comparison of the genotype-local ancestry (LA) correlation and genotype-PC correlation in NIGMS dataset. The distribution for LA is skewed to the right (or higher values of the correlation), indicating that multi-collinearity and thus inflated variance of estimated SNP effect size on gene expression, as quantified by the variance inflation factor of the ancestry predictor, *VIF*(*ancestry predictor*) = 1/(1 – *R*^2^)), is a greater problem for LA than for PC.

Using the NIGMS dataset, we showed that the genotype-local ancestry correlation was significantly higher than the genotype-PC correlation (one-sided Wilcoxon rank sum test, p-value<2.2×10^-16^, **Figure 1C**). This correlation can lead to inflated variance in the estimated *β*_*g*_, by the presence of the local ancestry *l* or the global ancestry *PC* in equation (1), as quantified by *VIF*(*γ*). We identified three SNPs with *VIF* of local ancestry larger than 10 (rs1314014, rs13313624 and rs186332) but no SNPs from the same *VIF* threshold for PC. This suggests that multi-collinearity is a greater problem for local ancestry than for PC and the confounding due to local ancestry is more likely to happen.

### Comparison of type I error rate and statistical power

Here we performed simulations to compare the effects of global ancestry and local ancestry adjustment for population structure on the type I error rate (**Table 1**). We used the actual genotypes of 81 NIGMS AAs and simulated gene expressions to be associated with the first PC or the average local ancestry of tested genes (see **Methods**). When the stratification is due to local ancestry (for example, effect size is the maximum of its distribution), the false positive rate is higher in the global ancestry adjustment than in the local ancestry adjustment (4.12×10^-3^>1.03×10^-4^). The inflation with no adjustment is larger when the stratification is due to local ancestry versus global ancestry (for example, effect size at the 100th percentile, 1.07×10^-2^>2.75×10^-4^). As expected, the inflation decreases as the effect size decreases. Importantly, adjusting for global ancestry was insufficient to remove stratification which may vary at each marker.

**Table 1.**
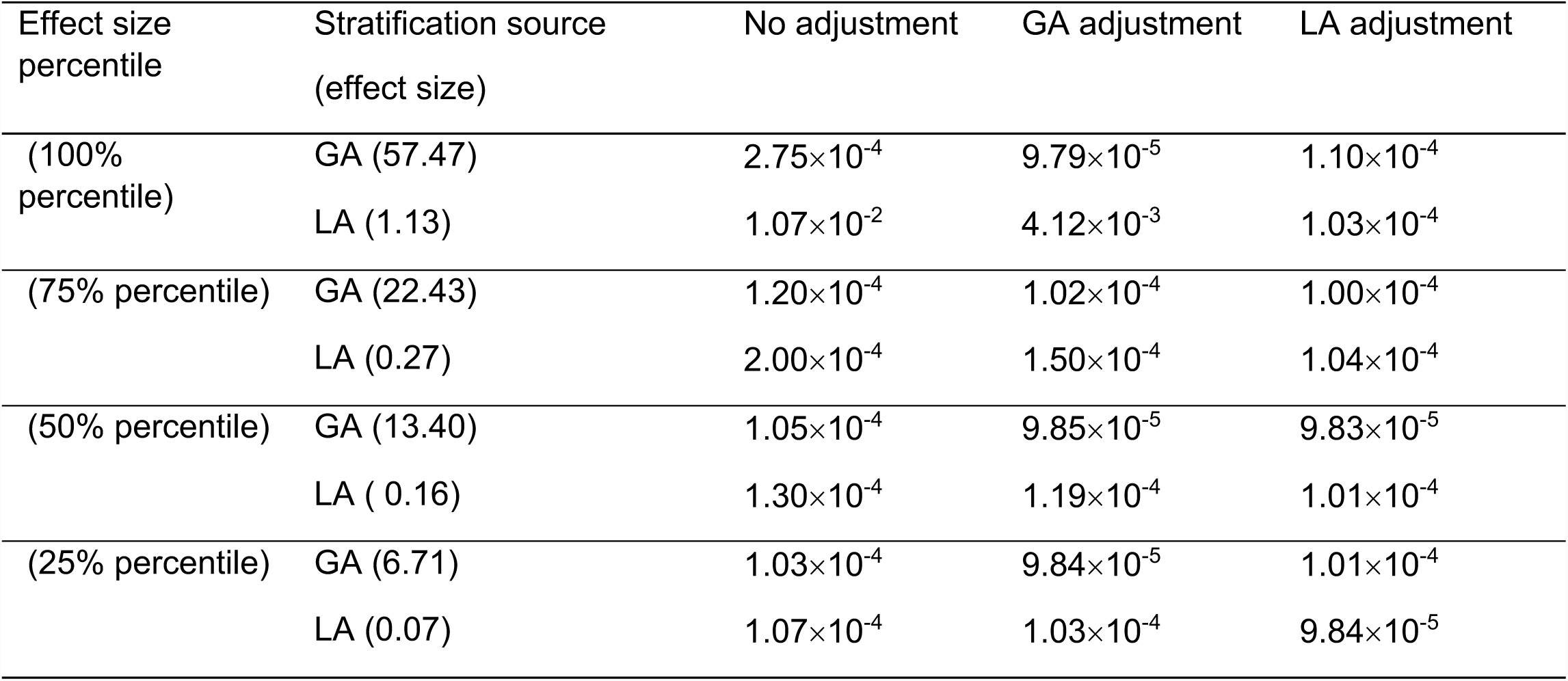
Type I error rate from analysis with no adjustment, global ancestry (GA) adjustment and local ancestry (LA) adjustment for population structure (false positives were identified using p<1×10^-4^)

| Effect size percentile | Stratification source (effect size) | No adjustment | GA adjustment | LA adjustment |
| --- | --- | --- | --- | --- |
| (100% percentile) | GA (57.47) | $2.75 \times 10^{-4}$ | $9.79 \times 10^{-5}$ | $1.10 \times 10^{-4}$ |
| | LA (1.13) | $1.07 \times 10^{-2}$ | $4.12 \times 10^{-3}$ | $1.03 \times 10^{-4}$ |
| (75% percentile) | GA (22.43) | $1.20 \times 10^{-4}$ | $1.02 \times 10^{-4}$ | $1.00 \times 10^{-4}$ |
| | LA (0.27) | $2.00 \times 10^{-4}$ | $1.50 \times 10^{-4}$ | $1.04 \times 10^{-4}$ |
| (50% percentile) | GA (13.40) | $1.05 \times 10^{-4}$ | $9.85 \times 10^{-5}$ | $9.83 \times 10^{-5}$ |
| | LA ( 0.16) | $1.30 \times 10^{-4}$ | $1.19 \times 10^{-4}$ | $1.01 \times 10^{-4}$ |
| (25% percentile) | GA (6.71) | $1.03 \times 10^{-4}$ | $9.84 \times 10^{-5}$ | $1.01 \times 10^{-4}$ |
| | LA (0.07) | $1.07 \times 10^{-4}$ | $1.03 \times 10^{-4}$ | $9.84 \times 10^{-5}$ |

To compare the type II error rate, we again used the actual genotype data and randomly selected SNPs to be causal eQTLs for pre-specified genes. When the gene expression was associated only with the genotype, the areas under the Receiver Operating Characteristic (ROC) curve for the identification of true eQTLs were similar between the two adjustment methods (**Figure 2A, C**, Paired *t*-test of area under the curve (AUC) for a false positive rate in the range 0-0.2 over 100 simulations: p-value=0.16). However, when the gene expression was associated with the SNP and its corresponding local ancestry simultaneously, the ROC curve for local ancestry adjustment was above that for global ancestry adjustment (**Figure 2B, D**, Paired *t*-test of AUC for a false positive rate in the range 0-0.2 over 100 simulations: p-value=1.03×10^-8^). Power comparison results show that the two adjustment approaches are equally powerful to identify true eQTLs, while local ancestry adjustment can substantially increase the power when gene expression changes with local ancestry.

**Figure 2.**
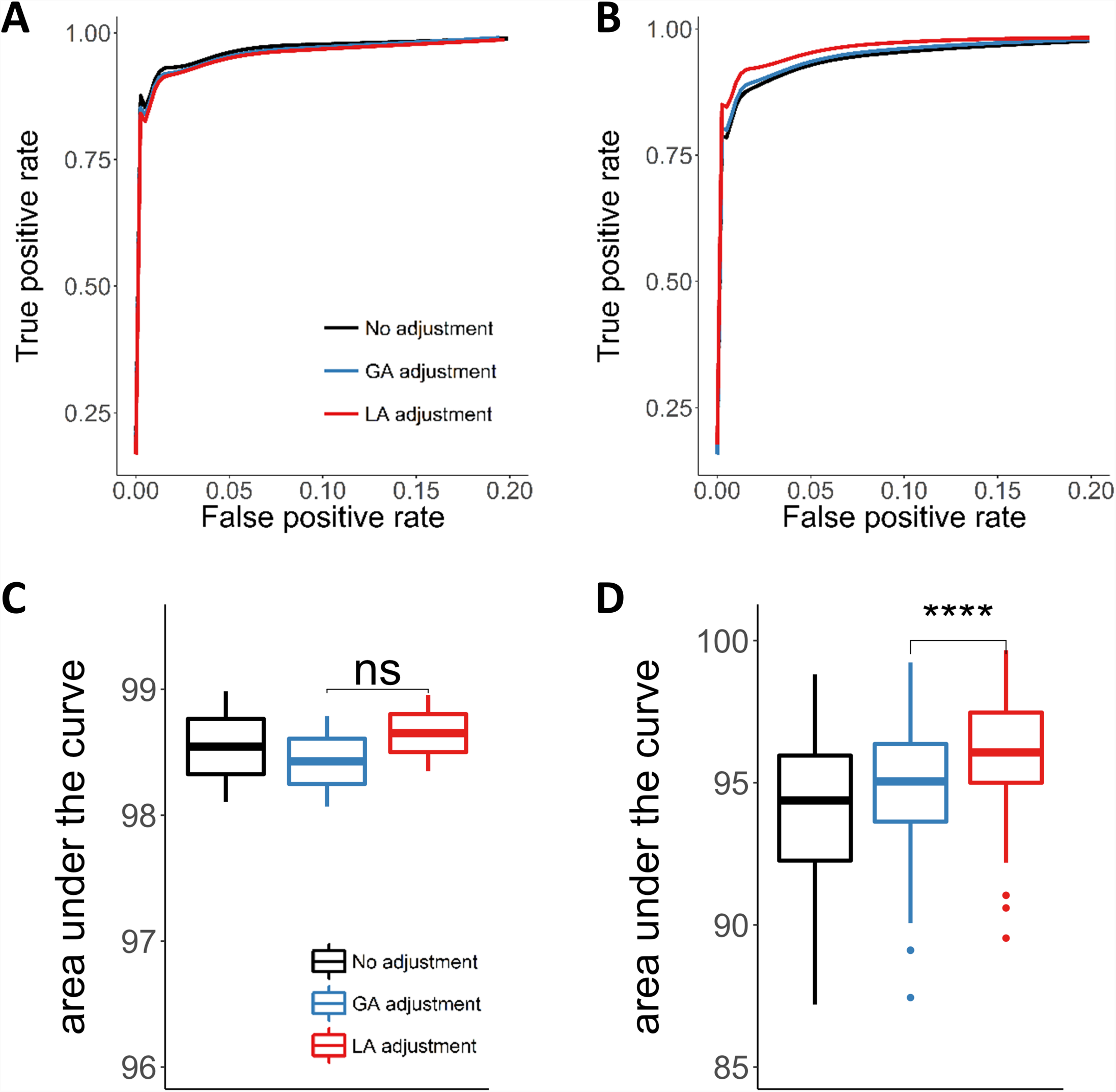
Power analysis for eQTL mapping with simulated data based on NIGMS dataset. We simulated the expression of 500 genes and calculated the association with a random sampling of 1000 SNPs using different methods to control for population structure confounding. Among 500,000 associations, we selected 50 SNPs to be true eQTLs. **A, C.** ROC curve (**2A**) and average AUC (**2C**) across 100 simulations when gene expression was associated with the SNP, showing equal performance (significance was calculated from a paired two-sided t-test). **B, D.** ROC curve (**2B**) and average AUC (**2D**) across 100 simulations when gene expression was associated with both SNP and local ancestry (LA), showing improved performance with LA adjustment.

### eQTL mapping in admixed samples

We developed an efficient approach, Joint-GaLA-QTLM, to eQTL mapping in a recently admixed population using local ancestry (see **Methods**). We applied Joint-GaLA-QTLM to cis-eQTL mapping in the NIGMS dataset. We adjusted for the top 3 PCs in the global ancestry adjustment method, and adjusted for the corresponding local ancestry of each tested SNP in the local ancestry adjustment method. We used a hierarchical correction method to select significant eQTLs (see **Methods**). We detected 270 eQTLs using the global ancestry adjustment method and 277 eQTLs using the local ancestry adjustment method. Among these eQTLs, 197 were shared by these two methods, while 21 and 14 eQTLs were detected only with the local ancestry and global ancestry adjustment, respectively. We compared the nominal (SNP association) p-values from the various methods (**Figure 3A**). The eQTLs found by both methods were more significant than the eQTLs unique to either method alone, suggesting that both methods were powerful to identify significant eQTLs.

**Figure 3.**
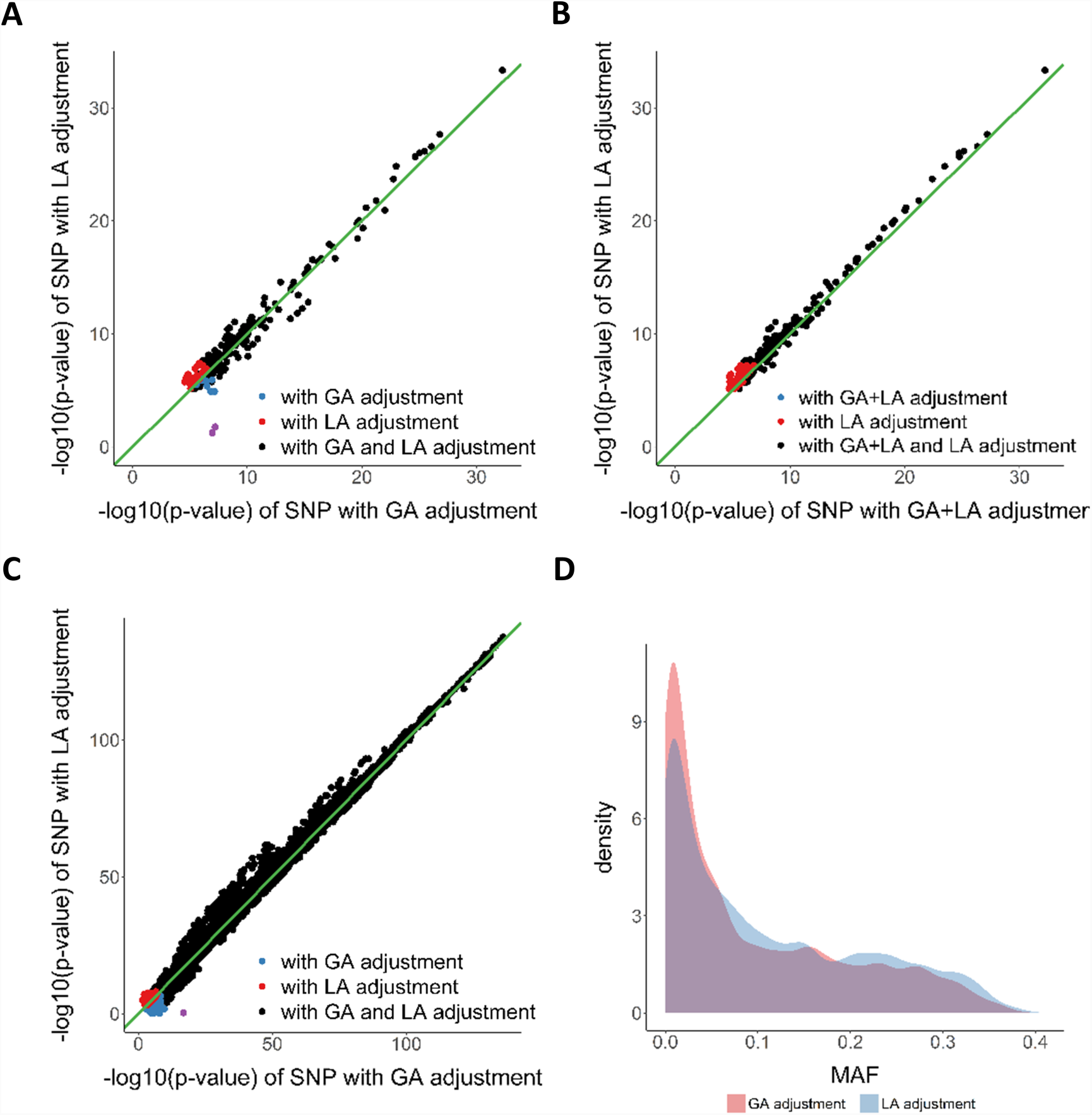
Comparison of eQTL mapping conducted using different ancestry adjustment methods. We performed eQTL mapping using global ancestry (GA) and local ancestry (LA) adjustment in the NIGMS dataset of AAs and the GTEx whole blood dataset (including EAs and AAs). The NIGMS is a recently admixed sample set while GTEx is a multiethnic sample set, and we sought to compare the approaches in both scenarios. Purple dots in A and C represent eQTLs, whose effect sizes were highly inflated in GA adjustment method and were found to be potential false positives. **A.** eQTL nominal p-values with GA adjustment or LA adjustment in NIGMS dataset, showing potential false positives (purple dots, **Figure S1**). **B.** eQTL nominal p-values with GA+LA adjustment or LA adjustment in NIGMS dataset, showing that LA adjustment alone (i.e., without the additional adjustment for global ancestry) may suffice. **C.** eQTL nominal p-values with GA adjustment or LA adjustment in GTEx whole blood dataset, showing potential false positive (purple dot). **D.** MAF distribution of eQTLs unique to GA or LA adjustment in GTEx whole blood dataset, showing a higher proportion of low frequency variants unique to GA adjustment.

We further investigated the eQTLs identified only by one method. Most of these method-specific eQTLs clustered at the margin of statistical significance. However, two eQTLs (rs8044834 with *AMFR*, p-value with global ancestry adjustment: 6.35×10^-8^, p-value with local ancestry adjustment: 1.74×10^-2^; rs2341000 with *PLA2G4C*, p-value with global ancestry adjustment: 1.12×10^-7^, p-value with local ancestry adjustment: 5.81×10^-2^) were highly significant only with global ancestry adjustment (**Figure 3A, Table 2**). Notably, local ancestry was significantly associated with gene expression at these loci (**Table 2**) and identified SNPs show large differentiation in allele frequency between CEU and YRI; thus we hypothesized that local ancestry confounded the eQTL association, resulting in false positive eQTLs. To test the hypothesis, we evaluated the association between genotype and gene expression in a subsample with two African ancestry alleles and in a HapMap CEU cohort (n=60), and found that the eQTL associations were no longer significant (Additional file 1: **Figure S1**). These eQTLs were not significant in GTEx (v7) LCL eQTL database as well. This highlights the possibility for spurious association between genotype and gene expression in loci where local ancestry is associated with gene expression. Among 21 eQTLs unique to local ancestry adjustment, 9 were significant (42.86%) in GTEx LCL eQTL database. For 14 eQTLs that were unique to GA adjustment, only 1 eQTL was significant (7.14%). The replication rate is significantly higher for eQTLs unique to local ancestry adjustment than for eQTLs unique to global ancestry adjustment (chi-squared test p-value=2.20×10^-2^).

**Table 2.** eQTLs unique to PC adjustment Two eQTLs (row 1, 2) were found to have highly significant associations by the global ancestry (GA) adjustment method, but were non-significant using the local ancestry (LA) adjustment method in NIGMS dataset (purple dots in **Figure 7A**); similarly, one such eQTL (row 3) was found in GTEx whole blood dataset (purple dot in **Figure 7C**). Included in the table are the p-values of allelic association tests with no correction for ancestry, with PC correction, with LA correction, p-value of LA in the allelic association with LA adjustment. Allele frequencies are from 1000 Genome Phase3 data.

| SNP<br>ref/alt | Gene | eQTL p-value,<br>no correction | eQTL p-value,<br>PC correction | eQTL p-value,<br>LA correction | LA<br>association<br>p-value | Alt allele<br>frequency in<br>YRI | Alt allele<br>frequency<br>in CEU |
| --- | --- | --- | --- | --- | --- | --- | --- |
| rs8044834<br>C/T | *AMFR* | $2.26 \times 10^{-9}$ | $6.35 \times 10^{-8}$ | $1.74 \times 10^{-2}$ | $8.80 \times 10^{-4}$ | 4% | 58% |
| rs2341000<br>G/T | *PLA2G4C* | $3.01 \times 10^{-8}$ | $1.12 \times 10^{-7}$ | $5.81 \times 10^{-2}$ | $7.17 \times 10^{-3}$ | 100% | 46% |
| rs2814778<br>C/T | *DARC* | $2.43 \times 10^{-44}$ | $1.67 \times 10^{-17}$ | $4.77 \times 10^{-1}$ | $1.99 \times 10^{-44}$ | 0% | 99% |

We then compared the results from local ancestry adjustment with results from alternative methods. We tested the effects of local ancestry plus global ancestry adjustment on the cis-eQTL mapping (**Figure 3B**). Surprisingly, the p-values with both adjustments were less significant than those with the local ancestry adjustment alone for shared eQTLs (Wilcoxon signed rank test: p-value <2.2×10^-16^), suggesting that including PC as additional adjustment for population structure to local ancestry will reduce power. By using only 1 or 2 PCs, we observed a similar pattern as the results based on 3 PCs (Additional file 1: **Figure S2**). We also ran the cis-eQTL analysis using the LMM approach (implemented in GEMMA) to control for population structure and cryptic relatedness. The GEMMA approach demonstrated higher statistical power compared to local ancestry adjustment but failed to remove the false positives (Additional file 1: **Figure S2**).

### eQTL mapping in multiethnic samples

Mapping eQTLs in GTEx data allowed us to evaluate the generalizability of our findings on the importance of local ancestry adjustment in a recently admixed population to multiethnic eQTL studies consisting of *both* subjects of relatively homogeneous ancestry and individuals of recent admixture. We applied Joint-GaLA-QTLM to GTEx LCL (n=114) and whole blood (n=356) datasets. Consistent with the results in the NIGMS dataset, more eQTLs were identified with local ancestry adjustment than with global ancestry adjustment (see Additional file 1: **Table S1**). Nominal p-values from local ancestry adjustment were more significant than those from global ancestry adjustment (Wilcoxon signed rank test; LCL dataset: p-value<2.2×10^-16^; whole blood dataset: p-value<2.2×10^-16^, **Figure 3C**) for shared eQTLs. In the whole blood dataset, we identified one SNP that was highly significant only with global ancestry adjustment (rs2814778 with *DARC*, p-value with global ancestry adjustment: 1.67×10^-17^, p-value with local ancestry adjustment: 4.77×10^-1^) and found its local ancestry and genotype had perfect correlation, again suggesting potential local ancestry confounding. Notably, eQTLs unique to global ancestry adjustment were more likely to have small MAF (MAF<0.10, chi-squared test: p-value<2.2×10^- 16^, **Figure 3D**) than those unique to local ancestry adjustment. Taken together, these results demonstrate the importance of local ancestry adjustment for cis-eQTL mapping even in samples with a relatively small proportion of admixture.

### Empirical study of PVE by local ancestry

We quantified the variance explained by local ancestry using a LMM model, which models a random effect according to the local admixture relatedness between individuals (see **Methods**). We estimated the distribution of PVE_*l*_ (mean=0.30, variance=0.08) in the GTEx muscle dataset AA samples (**Figure 4A**, Additional file 1: **Table S2**). The range of reliably estimated PVE_*l*_ (FDR<0.10) was [0.23, 0.99]. Genes with reliable PVE_*l*_ estimates were significantly enriched for differentially expressed genes (see **Methods**) between AAs and EAs (hypergeometric test: p=2.21×10^-6^), suggesting that PVE_*l*_ could be capturing the degree of population differentiation at causal variants, as also implied by our statistical model. Furthermore, the proportion (0.22) of genes with nominally significant PVE_*l*_ estimates (p<0.05) was much greater than expected by chance (0.05). The greater proportion of genes with significant PVE_*l*_ estimates than PVE_*g*_ (Additional file 1: **Table S2**) raises the possibility that joint analysis of local ancestry and genetic variation may improve heritability estimation in this population.

**Figure 4.**
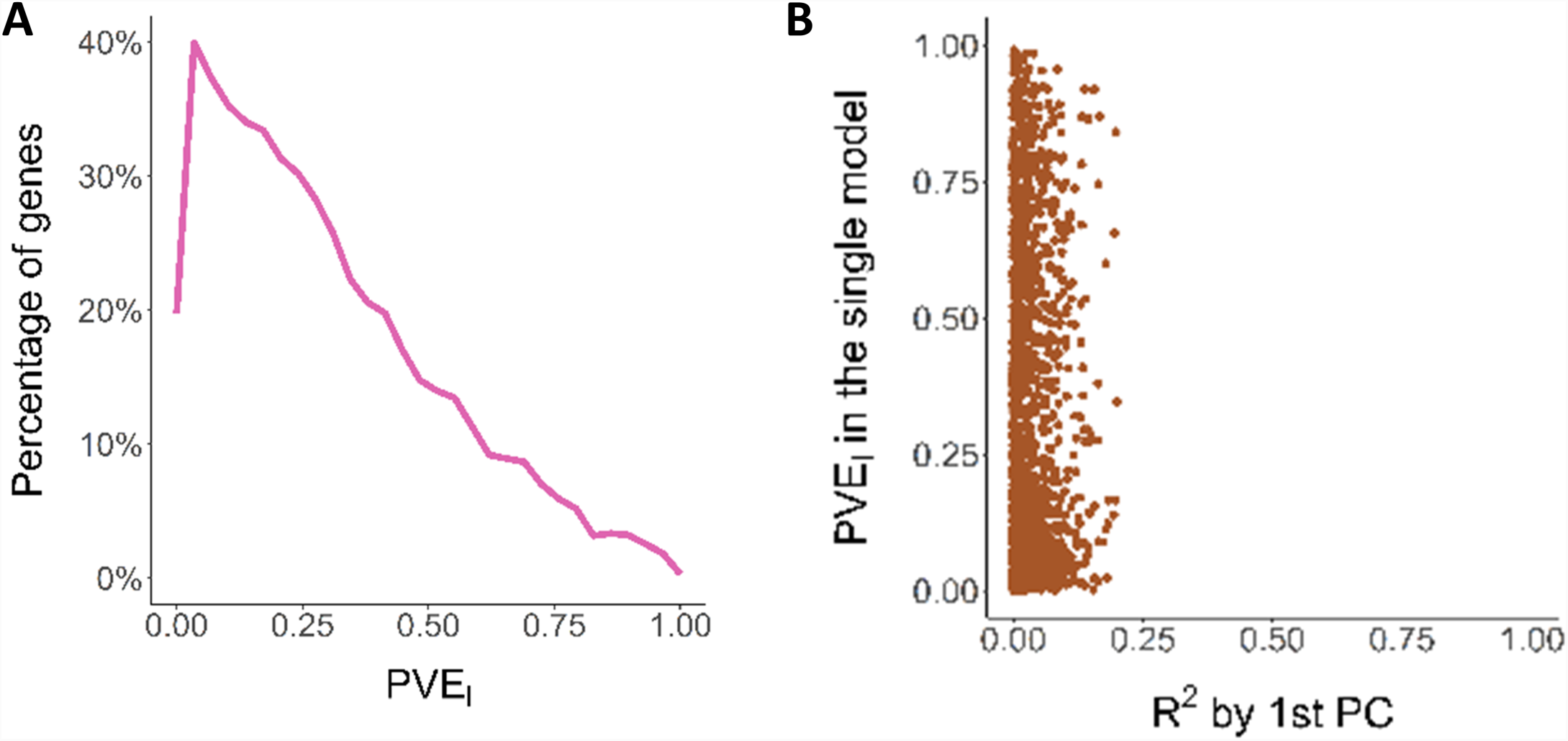
PVE_*l*_ analysis in AAs. To determine the variance explained by local ancestry, we estimated PVE_*l*_ for genes in the GTEx skeletal muscle dataset in AAs. The *R*^2^ of global ancestry (PC1) from simple linear regression with gene expression did not capture the variance 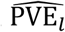 explained by local ancestry. **A.** Distribution of 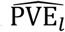 (total of 1,740 gene). **B.** Comparison of 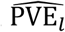 and *R*^2^ of global ancestry (PC1) from simple linear regression with gene expression.

When overlapping genes with reliable PVE_*l*_ estimates with the same number of genes selected according to the significance of the association between gene expression and the first PC, we found no shared genes, indicating the extent to which the global ancestry failed to capture the variance explained by the local admixture structure. In GTEx muscle data, *R*^2^ from linear regression of the global ancestry (PC1) with gene expression tended to underestimate the variance PVE_*l*_ explained by local ancestry (**Figure 4B**). When we used the local ancestry from the entire genome to construct the genetic relatedness matrix, we identified no genes with reliable estimates (FDR<0.10), suggesting either that the gene expression variation was more related to the local, instead of the global, admixture structure or that we were underpowered to obtain a precise estimate (in analogy with estimating the trans-eQTL contribution using a trans-eQTL-based GRM).

### Simulation studies of heritability estimation in an admixed population

We designed extensive simulations, using real and simulated (admixed) genotype data, of diverse genetic architectures (see **Methods**), to compare three methods for estimating the heritability of gene expression in admixed populations. The first method, Simple-LMM, applies restricted maximum likelihood (REML) to obtain an estimate. The second method, LDSR [21], estimates the confounding due to population stratification (from the “intercept”) as well as the trait heritability (from the “slope”) by regressing the GWAS test statistics on LD scores. We propose a novel method, Joint-GaLA, which includes a local ancestry component when estimating the heritability. In all three methods, we control for global ancestry (PC1) to remove potential confounding due to global ancestry.

In simulations with simulated genotype data, the PVE_*l*_ estimates derived from REML were in line with equation (2*) analytically derived from the statistical model (Additional file 1: **Table S3**), confirming the expression for the estimate 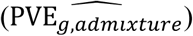 of the expected heritability in the presence of admixture (see **Methods**). From equation (2*), the trait variance explained by local ancestry (PVE_*l*_) may therefore be reflecting the fixation index 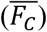 at causal variants and/or “tagging” of causal variant effect (PVE_*g*_) on phenotype. Furthermore, the Gaussian approach, versus the (more computationally intensive) mixture model approach, to modeling the effect explained by local ancestry (see **Methods**) was sufficient to provide accurate estimates (Additional file 1: **Table S3**), consistently across all choices for the number of causal variants.

We simulated gene expression with local ancestry effect (several percentiles chosen from real data to represent different degrees of stratification). 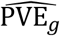 estimates from Simple-LMM and LDSR, with control only for global ancestry, tended to suffer from upward bias (**Figure 5A**) whose magnitude increased with a greater degree of stratification, across the range of number of causal variants tested (Additional file 1: **Figure S3)**. In all cases, Joint-GaLA was closer to the assumed heritability and significantly different from Simple-LMM (median values, using 10, 25, 100, 200, and 500 causal variants, of (0.492, 0.507, 0.500, 0.508, 0.498) vs (0.366, 0.351, 0.309, 0.335, 0.286) for Simple-LMM and Joint-GaLA when PVE_*l*_=0.2 and PVE_*g*_=0.3, respectively; Mann-Whitney U test p<0.002 for all comparisons between the two methods). Estimates from LDSR showed a significantly larger standard error than Joint-GaLA (Mann-Whitney U test p=0.008).

**Figure 5.**
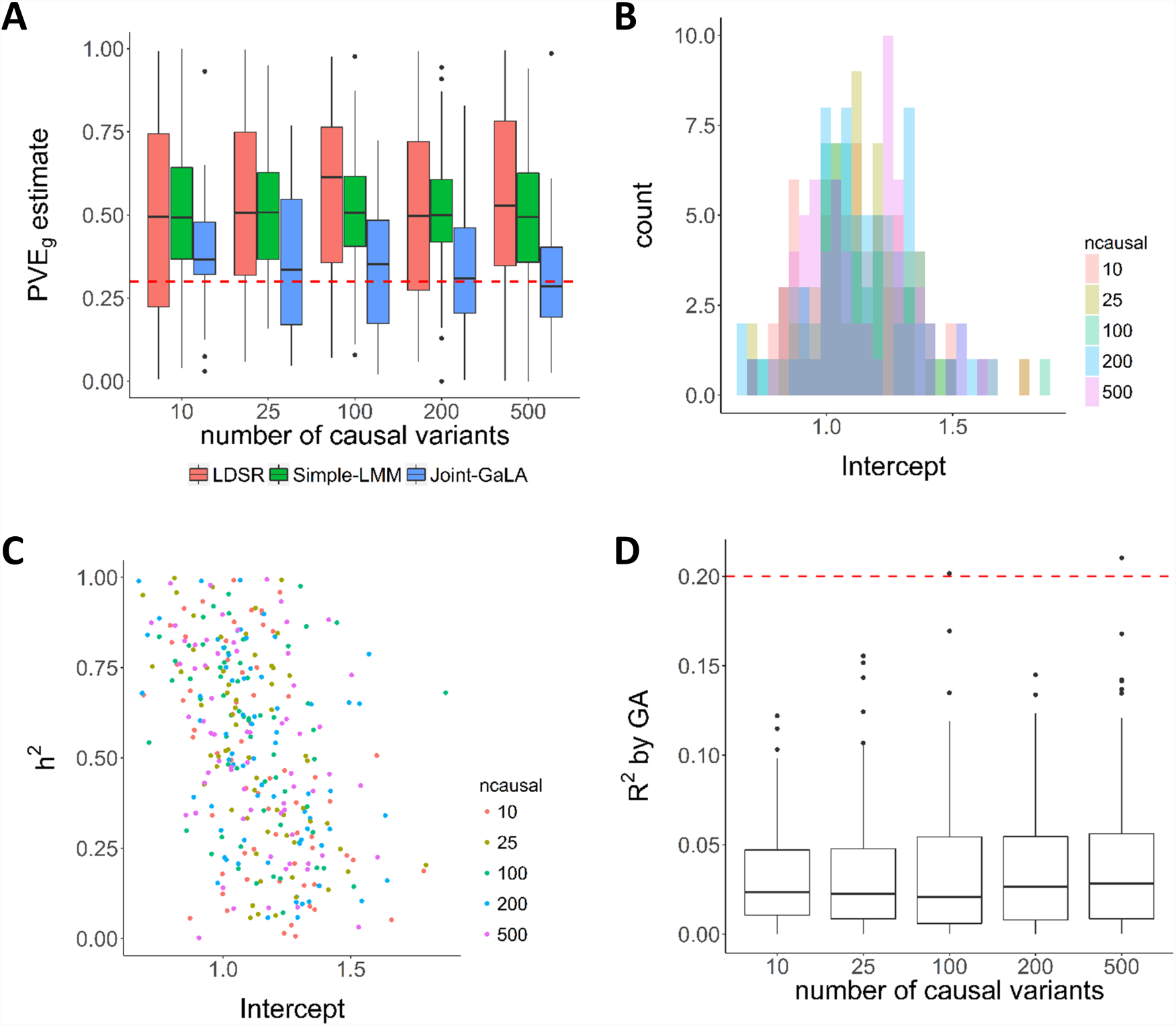
PVE_*g*_ estimation in simulations. We performed simulations with real genotype data to evaluate the accuracy of heritability estimation with Simple-LMM, Joint-GaLA, and LDSR. We assumed the effect of local ancestry on gene expression (PVE_*l*_) was 0.2 (one of several levels of stratification tested, based on empirical data) and varied the number of causal variants. **A.** The assumed heritability was 0.30, shown as a dashed horizontal line. The estimates 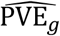 from Simple-LMM were substantially inflated in comparison with those from Joint-GaLA and nearly identifical to those from LDSR, across the range of number of causal variants tested. LDSR showed the widest variation in the estimates. Joint-GaLA was closer to the expected heritability than Simple-LMM and showed significantly improved estimates for all comparisons (based on number of causal variants; Mann-Whitney U test p<0.002). **B.** The intercept estimates from LDSR assuming the same genotype data and a fixed local ancestry effect on phenotype showed wide variation. **C.** The estimates for heritability were negatively correlated with the intercept estimates for the amount of stratification. Note the presence of inflated estimates of heritability observed even under low estimated levels of confounding (e.g., near 1 for the intercept). **D.** *R*^2^ from global ancestry (PC1), estimated from linear regression, substantially underestimated the trait variance explained by local ancestry, across all choices for the number of causal variants. The dashed line shows the expected trait variance explained by local ancestry.

Simple-LMM and LDSR generally gave near-equivalent estimates of heritability, across the range of number of causal variants tested (**Figure 5A**), but LDSR estimates had substantially larger variability (Mann-Whitney U test p=0.008). As an estimate of population confounding, the intercept from LDSR showed wide variation (**Figure 5B**), with a higher estimate of population confounding associated with greater uncertainty (i.e. larger standard error) in the estimate (Spearman’s rho=0.555, p<2.2e-16). The estimates for heritability were negatively correlated (Spearman’s rho=-0.45, p<2.2e-16) with the intercept estimates for the amount of confounding, with inflated estimates of heritability observed even under low estimated levels of confounding (**Figure 5C**). Note that LDSR, by design, does not provide an estimate for PVE_*l*_, and because of the wide variation in the estimate for the intercept, we would caution against using the intercept as a proxy for population confounding due to local ancestry.

Furthermore, we found that the *R*^2^ from global ancestry, estimated from linear regression, substantially underestimated the trait variance explained by local ancestry, across all choices for the number of causal variants (**Figure 5D**).

Simulating gene expression data (n=100) as the sum of causal genetic variation and an environmental component that is correlated with global ancestry (informed by median *R*^*2*^=0.009 between PC1 and gene expression from real data) with simulated genotyped and local ancestry data and assuming sparsity (n=10 causal variants) and polygenicity (n=1000 causal variants) and true heritability of 0.30, Joint-GaLA provided an unbiased estimate of heritability (median=0.307, mean=0.325, mean(SE)=0.052; median=0.300, mean=0.294, mean(SE)=0.098, respectively).

Finally, in simulations with real genotype data and with stratification due to local ancestry, we observed significant departure from the expected heritability 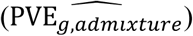 (see **Methods**; Mann-Whitney U test p<2.2e-16).

### Empirical study of PVE by genetic variation in admixed population

We utilized the GTEx skeletal muscle data to gain further insights into PVE_*g*_ (see **Methods**) in the largest RNA-Seq and whole genome sequencing data of AAs available to us. Using Simple-LMM, we estimated the distribution of PVE_*g*_ in the AA samples (mean=0.30, variance=0.05) and in the EA samples (mean=0.25, variance=0.04) of the same sample size (**Figure 6A**). Additional file 1: **Table S2** contains summary data on the estimates in the two populations in this tissue and Additional file 2 contains all PVE_*g*_ estimates. We identified genes with nominally significant PVE_*g*_ estimates (defined as p-value<0.05) in one population but not in the other, suggesting “*population-specific regulation.*” The comparison of PVE_*g*_ for genes with nominally significant estimates in both populations showed a modest but significant correlation (Spearman’s ρ=0.33, p-value=1.28×10^-7^; **Figure 6B**). At a more stringent threshold (false discovery rate (FDR)<0.10), we continued to observe a significant correlation (Spearman’s ρ=0.44, p-value=0.01; Additional file 1: **Figure S4**). We found no significant correlation between 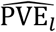 and 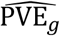 (Spearman’s ρ=0.04, p-value=0.21; **Figure 6C**) in AA samples.

**Figure 6.**
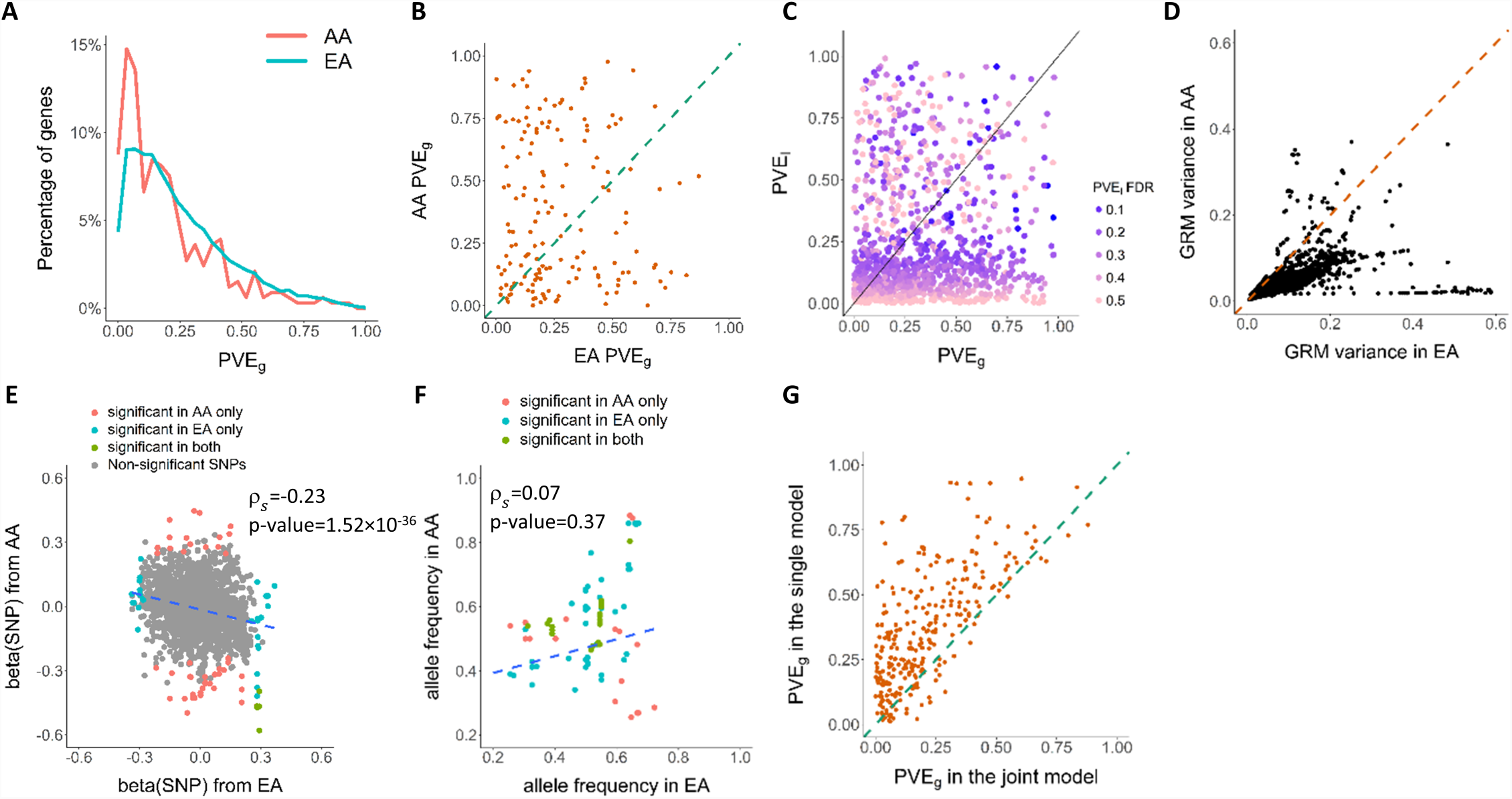
PVE_*g*_ analysis in EAs and AAs. We estimated the PVE_*g*_ for gene expression traits in the GTEx skeletal muscle dataset for African Americans (AAs) and an equal sample size (n=57) of Europeans (EAs) separately. Although there was a significant correlation in 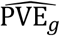 between the populations, many genes with nominally significant estimates (p-value<0.05) were discordant between the populations (**6B**). We investigated the contribution of variance in genetic relatedness (**6D**), effect size (**6E**) and allele frequency (**6F**) to the population specificity of PVE_*g*_. Comparison of 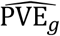 and 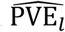 showed low correlation across the genes. We then fitted, for each gene, a joint model (Joint-GaLA) consisting of both genetic variation and local ancestry, to estimate the change in the estimate for PVE_*g*_. Most genes showed decreased estimate for PVE_*g*_ with the incorporation of local ancestry in the model, suggesting that local ancestry may explain some of the gene expression variation. **A.** Distribution of 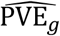 in AAs and EAs (total of 8,832 and 8,670 genes in AAs and EAs, respectively). **B.** Comparison of 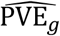 among genes with nominally significant estimates (p-value<0.05) in both AAs and EAs (total of 253 genes) showing a significant correlation. Similar result is observed at FDR<0.1 (Additional file 1: **Figure S2**). **C.** Comparison of 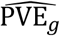 and 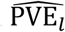 (points color-coded according to the FDR for 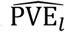). **D.** Comparison of the variance of the local genetic relatedness between AAs and EAs for all 19,850 genes, with EA showing significantly greater variance (one-sided Wilcoxon signed rank test, p-value<2.2×10^-16^). **E.** Example of a gene, *ZCCHC24*, for which local SNPs have opposite allelic direction between EAs and AAs. (The gene is not differentially expressed between the two populations.) Spearman correlation for all SNPs and the corresponding p-value are shown, and blue dashed line is fitted regression line. **F.** Example of a gene, *DDT*, for which SNPs associated with expression level (nominal p-value<0.05 from LMM association in either population) are population differentiated in allele frequency. Spearman correlation and p-value are shown, and blue dashed line is fitted regression line. **G.** Comparison of 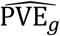 between Simple-LMM and Joint-GaLA.

We investigated the possible sources of the imperfect correlation in the estimated PVE_*g*_. The variance in genetic relatedness can be written as the sum of LD correlation over all pairs of SNPs used to construct the genetic relatedness matrix (GRM) (see **Methods**) [25]. The difference between the two populations in local LD pattern near each gene, estimated using the variance in the GRM (**Figure 6D**), can influence the estimated standard error of PVE_*g*_. We provide two examples to illustrate additional reasons for the population difference. LMM association analysis of the gene *ZCCHC24* 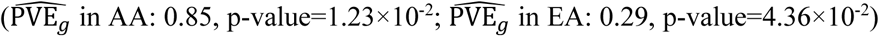 showed the effect sizes of local SNPs were negatively correlated between the two populations (Spearman’s ρ=-0.23 p-value=1.52×10^-36^; **Figure 6E**), suggesting population-dependent regulation with an “*opposite allelic direction*.” We compared the allele frequency of SNPs associated with *DDT* expression (nominal p-value<0.05 in either population) between EAs and AAs (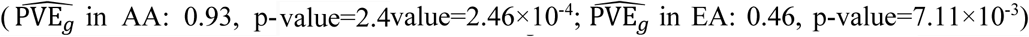 and found no evidence for correlation (Spearman’s ρ=0.07, p-value=0.37; **Figure 6F**). In both examples, although the gene had a significant PVE_*g*_ in both populations, the gene was nevertheless associated with a different set of variants (which were not in LD) in the different populations, suggesting “*alternative genetic regulation*”. For example, among the 50 SNPs that were associated with *ZCCHG24* expression in EAs, only 5 were in LD (LD>0.8) with associated SNPs in AAs. Finally, the polygenicity or sparsity of gene expression, which we explore in the next section, may differ for a given gene in the two populations.

We finally applied Joint-GaLA in the GTEx AA samples. Interestingly, we found that the PVE_*g*_ estimates from the Simple-LMM model (see **Methods**) tended to be inflated in comparison with the PVE_*g*_ estimates from the Joint-GaLA model (**Figure 6G**). This is consistent with simulations, in which Joint-GaLA outperformed Simple-LMM across all choices of number of causal variants when local ancestry contributed to the variance in phenotype.

### Sparsity or polygenicity of gene expression in an admixed population

We sought to characterize the sparsity or polygenicity of gene expression traits in this admixed population and compared the results of the PVE analysis from the LMM approach (see **Methods**), which is suitable for infinitesimal genetic architectures, and from a Bayesian sparse linear mixed model (BSLMM, all estimates in Additional file 3), which assumes a mixture distribution of effect sizes. The two approaches were highly correlated in their estimate of the polygenic component (Spearman’s ρ=0.82 between BSLMM-derived 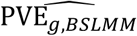 and LMM-derived 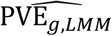, p-value<2.2×10^-16^) (Additional file 1: **Figure S5**). Nevertheless, we also identified genes for which BSLMM analysis showed a highly sparse local genetic architecture, i.e., genes with high estimated PGE (the proportion of gene expression variance explained by sparse genetic effects) and also high estimated PVE_*g*,*bslmm*_ (**Figure 7A**). Furthermore, the estimated total sparse genetic effect PGE was largely independent of the estimated total polygenic effect PVE_*g,lmm*_ across all genes tested as well as across all genes with a nominally significant estimate of PVE_*g,lmm*_ (p-value<0.05; **Figure 7B**).

**Figure 7.**
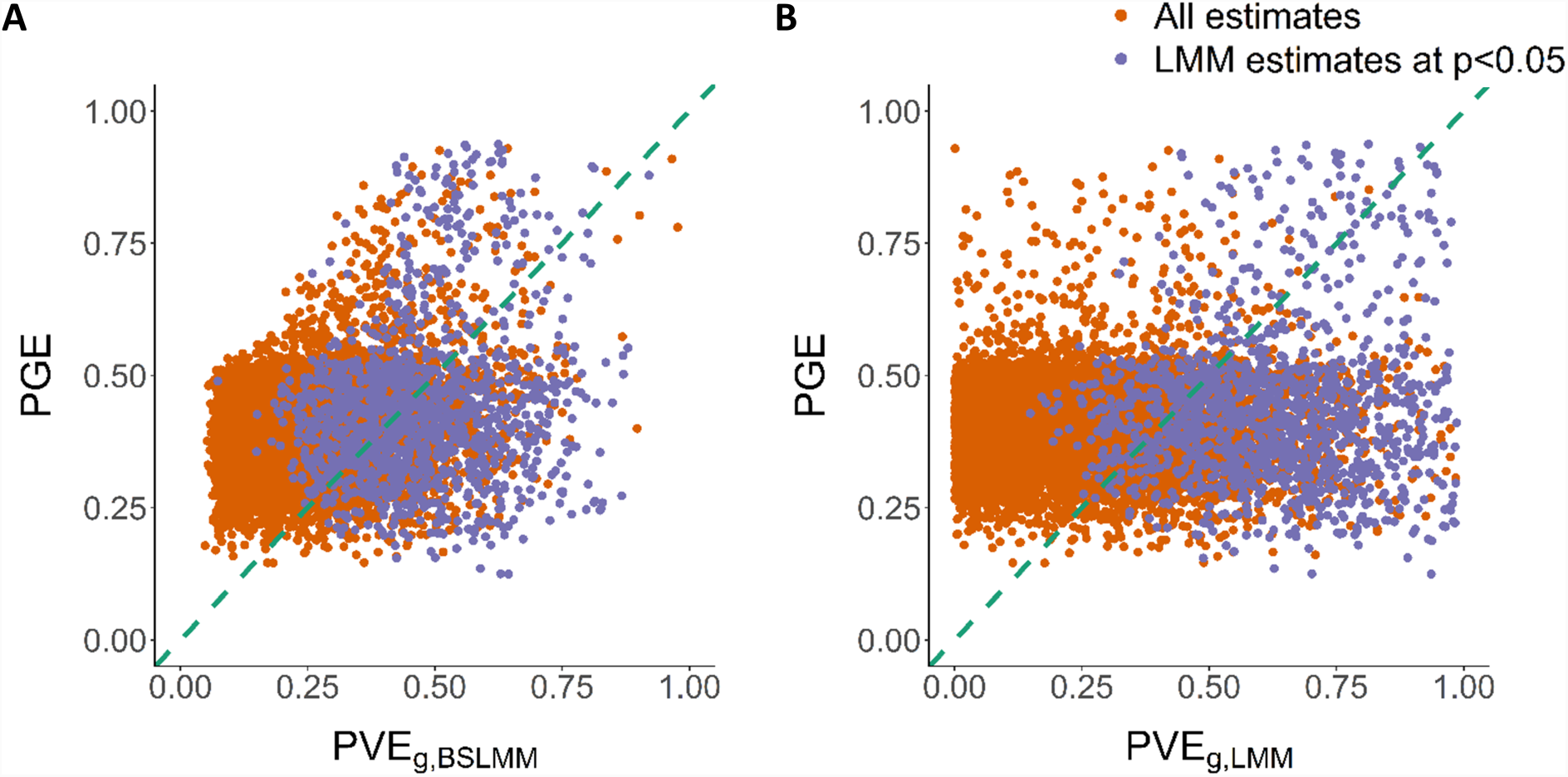
Sparsity and polygenicity of gene expression in AAs. We characterized the sparsity or polygenicity of gene expression traits using BSLMM analysis in AA GTEx skeletal muscle data. We estimated the proportion of variance in gene expression that can be explained by sparse effects (PGE) and the proportion of variance in gene expression that can be explained by sparse effects and random effects together (PVE_*g*,*BSLMM*_), which is most equivalent to our LMM-based PVE_*g*,*lmm*_. Genes whose estimated PVE_*g*,*lmm*_ was significant at p-value<0.05 were defined as nominally significant estimate A. The comparison of 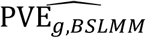 and PGE estimate from BSLMM. Genes with large 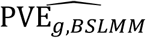 as well as large PGE estimate indicate a likely highly sparse local genetic architecture. B. The comparison of 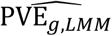 from GCTA and PGE estimate from BSLMM, showing the independence of the two components.

## Discussion

This study comprehensively evaluated the use of local ancestry in the analysis of genetic regulation of gene expression in an admixed population through extensive simulations and in real datasets. We developed a statistical model that allowed us to analytically formulate the relationships among global ancestry, the level of population differentiation at a causal eQTL, the trait variance explained by local ancestry, and the eQTL effect size. The model provides insights into potential sources of bias in the estimated regulatory effect of genetic variation on gene expression, including the degree of population differentiation and the uncertainty in local ancestry estimation. We extended this framework to the study of multiple causal eQTL variants. As a corollary of the model, characterization of gene expression in terms of sparsity or polygenicity has important implications for estimating the phenotypic variance explained by local or global ancestry. Hence, we comprehensively quantified the sparse genetic component and the polygenic component of gene expression in a recently admixed population, though this analysis was limited to a single tissue. Multi-tissue studies in a much larger sample size should facilitate additional insights into genetic architecture.

We performed a comprehensive analysis of the variance explained by local ancestry around each gene and across the genome to gene expression variation. In simulations with different degrees of stratification – informed by empirical data – due to local ancestry, an approach that incorporated local ancestry into heritability estimation (as in Joint-GaLA) provided a more accurate estimate of heritability in an admixed population than a naïve approach (as in Simple-LMM) that controlled only for global ancestry (e.g., as quantified by principal components). In these simulations, Simple-LMM and LDSR provided near-equivalent estimates of heritability. Both methods showed upward bias when controlling only for global ancestry in the presence of local ancestry stratification, although LDSR had significantly larger standard errors. Furthermore, the LDSR intercept, a measure of population confounding, showed wide variation, with a higher estimated level of confounding significantly associated with a greater degree of uncertainty in the estimate. Finally, under stratification, the estimated amount of confounding was found to be significantly (negatively) correlated with the estimated heritability in LDSR, indicating inflated estimates of heritability (slope) despite low reported levels of population confounding (intercept). As another corollary, the confounding can distort cross-population analyses of the contribution of genetic variants to gene expression variation. In particular, studies, in which one of the populations is admixed, that investigate the population specificity or sharedness of regulatory effects without taking into account local ancestry may suffer from this confounding. With diminished assumed level of stratification due to local ancestry in simulated genotype data, Joint-GaLA and Simple-LMM approached near-identical estimates of heritability, and LDSR, given its equivalence with Simple-LMM, would facilitate more reliable heritability estimation in this more controlled context.

Applying PVE_*g*_ estimation to real data, we observed a modest but significant correlation in estimated overall genetic effect between the populations, suggesting the existence of “shared regulatory architecture” for a number of genes. We investigated several factors underlying the population specificity of PVE_*g*_. The standard error of the estimate is closely related to the LD structure; thus local ancestry transitions present challenges for PVE analysis in recently admixed populations. Indeed, as our statistical model implies, local ancestry transitions can contribute to population differences in the estimated PVE_*g*_. Furthermore, our study would suggest that PVE estimation methods that explicitly incorporate LD adjustment might yield larger power [26]. Given the small sample size, for nearly half of expressed genes (33.38% in AAs and 45.32% in EAs), we could not obtain PVE_*g*_ estimates because the phenotypic variance-covariance matrix is not positive definite. This observation demonstrates the necessity of large sample size for PVE analysis even for intermediate (e.g., molecular) phenotypes. We developed an R package, LAMatrix, which adjusts for local ancestry in eQTL mapping and implements Joint-GaLA-QTLM in a computationally efficient framework. Our implementation can be exploited in studies that incorporate a SNP-level covariate (e.g. epigenetic marker or structural variant), which may prove crucial in disentangling the influences of various factors on a cellular phenotype. We illustrated with simulations that type I and type II errors will be inflated when gene expression is associated with local ancestry, which was observed for a substantial number of genes in both admixed samples and multiethnic samples. The application of Joint-GaLA-QTLM to the NIGMS dataset (admixed) and GTEx whole blood/LCL dataset (multiethnic) showed that our approach displayed greater power to identify eQTLs than the prevailing approach that adjusts for global ancestry. In GTEx whole blood study, more eQTLs unique to GA adjustment have small MAF, which is vulnerable to false positives [27], again supporting that the proposed local ancestry adjustment is more powerful to identify true eQTLs.

Finally, our approach can be easily extended to the search for the genetic determinants of other high-dimensional omics phenotypes (for instance, to enlarge our understanding of the epigenome and the proteome) in more heterogeneous populations.

## Conclusions

We show that use of local ancestry can improve identification of regulatory variants (QTL mapping) and estimation of their total effect (heritability estimation), with broad implications for genetic studies of complex traits. Taken together, these results substantially extend existing approaches and provide a framework for future large-scale studies of genetic regulation of gene expression in multiethnic or admixed samples.

## Methods

### Genotype data

We downloaded GTEx v7 genotype data (635 individuals) from dbGaP (phs000424.v7.p2). The genotype data contain individuals with recent admixture (e.g., African Americans) [2] as well as individuals of more homogeneous (European) ancestry, the latter comprising the majority of the samples (∼85%). We performed the MAF>0.01 filtering following GTEx and removed all multiallelic SNPs and SNPs on the sex chromosomes. The number of SNPs left was 9,910,646. GTEx data were used for PVE analysis and eQTL mapping in multiethnic samples.

We used 100 AA samples that were part of the NIGMS Human Variation Panels to assess the impact of local ancestry in pure admixed populations. We downloaded the genotype intensity files from dbGaP (dbGaP study accession phs000211.v1.p1) The genotyping had been performed on Affymetrix Genome-Wide Human SNP Array 6.0 platform containing 908,194 SNPs. We used the Affmetrix Genotyping Console to process the genotype intensity files and to call the genotypes on the forward strand. We kept 83 individuals with gene expression measurements. We merged the genotype with 1000 Genome phase 3 genotype data and performed the PCA with PLINK [28]. Two individuals with partial East Asian ancestry were removed from the subsequent analysis, leaving 81 samples. Quality control was performed with PLINK. We removed SNPs that are on the sex chromosomes, have duplicated positions, are multiallelic in 1000 Genome reference panel, are out of Hardy-Weinberg equilibrium (p-values<1×10^-5^), and whose genotyping missing rate is larger than 5% and minor allele frequency (MAF) is less than 5%. The total number of SNPs remaining in the analysis after the quality control was 724,100. This dataset was used for simulation and eQTL mapping. We also used 60 CEU (U.S. residents with northern and western European ancestry) samples from Phase 2 HapMap (release 23) and kept 714,082 SNPs which were a subset of the NIGMS AA dataset. This dataset was used for the replication of eQTLs detected in the NIGMS AA dataset.

### Local ancestry estimation

After quality control, the genotype data were phased with SHAPEIT [29] using 1000 Genome phase 3 in build 37 coordinates as the reference genome. We utilized the YRI samples (Yoruba people of Ibadan, Nigeria) and CEU samples from the 1000 Genome Phase 3 as the reference ancestral genomes to estimate the local ancestry (0, 1 or 2 African ancestry alleles) using a conditional random field based approach, RFMix [12]. When performing local ancestry inference, RFMix models strand-flip errors to account for potential phase errors. The window size in RFMix was set to be 0.15 Mb for the GTEx data and 0.20 Mb for the NIGMS data because the latter have less SNPs. We compared the first PC with the average local ancestry across the genome; this comparison shows a nearly perfect correlation in both the NIGMS and GTEx datasets (Additional file 1: **Figure S6**), demonstrating reliable estimation of local ancestry.

### Gene expression data

We used the gene expression data from GTEx v7 skeletal muscle dataset (n=491 with genotype data, of which n=57 are AA samples) for PVE estimation (see “PVE estimation in real transcriptome data”). This tissue was used because it has the largest number of AA samples in the GTEx data. The expression values have been normalized for 19,850 autosomal genes. We used GTEx whole blood (n=369) and cell-EBV-transformed lymphocytes (LCL, n=117) datasets to test our eQTL mapping approach in a multiethnic population (see “Cis-eQTL mapping in NIGMS and GTEx”). There were 19,432 and 21,467 expressed autosomal genes in these two datasets, respectively.

We obtained gene expression data for 81 AAs (represented in the NIGMS dataset) and 60 HapMap CEU samples from Gene Expression Omnibus (GEO) with the accession number GSE10824 [30]. The expression intensity for 8,793 probles was quantile normalized and corrected for background with the Robust Multichip Average (RMA) method. We filtered probes whose variances were less than the 0.4 quantile of variances of all genes, probes without Entrez Gene ID, duplicated probes, and probes on sex chromosomes. We performed log_2_ transformation on the gene expression data. A total of 4,595 probes representing 4,595 genes were included in the analysis after the quality control. We converted the probe IDs to the gene symbols using the HG Focus annotation file and obtained gene positions from the GENCODE release 19.

### Statistical model

Let *i* denote the *i*-th individual and *f* denote a local causal genetic variant for a gene. Then gene expression can be written as follows [24]:

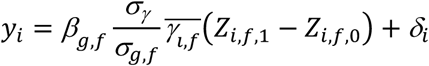

Here *β*_*g*,*f*_ is the effect size of the genetic variant *f* on gene expression trait y, 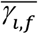 is the normalized local ancestry (*γ* – E[*γ*_*i,f*_])/ *σ*_*γ*_, *σ*_*γ*_ is the variance of local ancestry, *σ*_*g*,*f*_ is the variance of genotype at SNP *f*, and *δ*_*i*_ is the residual. *Z*_*i*,*f*,∗_ are Bernoulli distributed according to the allele frequency of the SNP *f* in population 0 (*p*_*f*,1-_) or 1 (*p*_*f*,0_),

### Single causal variant

*β*_*r*,*f*_, which is the effect explained by local ancestry at the SNP *f*, can be estimated from VAR [E[*y*_*i*_|*γ*_*i*,*f*_]]. If we assume a single causal eQTL variant, such as often assumed to simplify certain types of eQTL analysis [31, 32], we obtain the following:

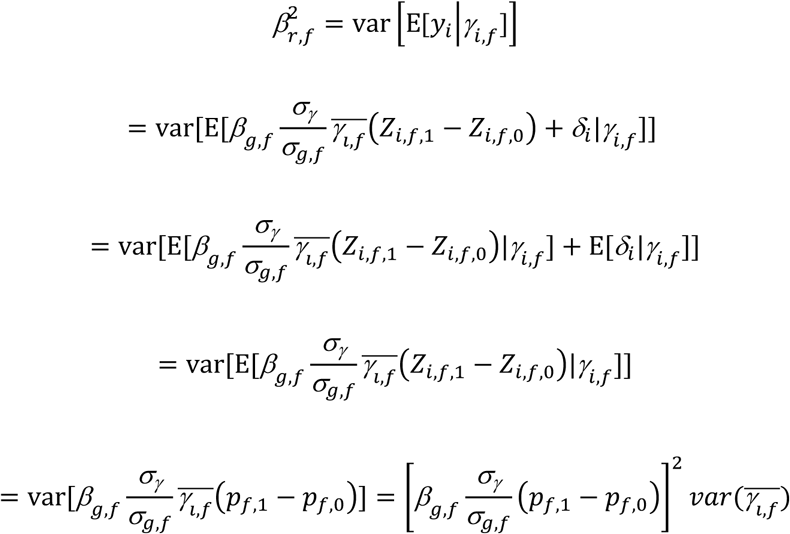

Since 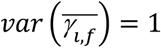 and 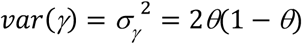 where *θ* is the global ancestry, we obtain:

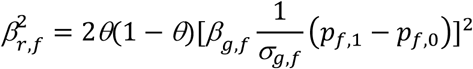

Let

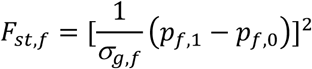

which is a measure of population differentiation or allele frequency difference at the variant *f*. Then the following expression, which relates the effect explained by local ancestry, global ancestry, the effect of the genetic variant, and the degree of population differentiation in a single equation, follows:

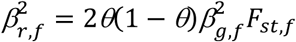

#### Multiple casual variants

We sought to generalize equation (1) to the case of multiple casual eQTL variants in the cis region. Here it matters for the purpose of estimating the variance PVE_*l*_ explained by local ancestry whether there is any local ancestry transition in the region and how many such transitions exist. (Local ancestry segments may extend over a large distance.) Suppose there are *n* local ancestry transitions. This implies 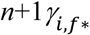 local ancestry classes in the region (with *f*\* being the local ancestry membership of the variant *f*). (A stretch of the genome in between local ancestry transitions represents a local ancestry class.) Let *m* be the number of local causal genetic variants for the expression of the gene. (In what follows, we will assume there are no other causal (e.g., trans) eQTLs outside the region, strictly restricting our focus on cis variants.) Then we obtain, following [24], the following:

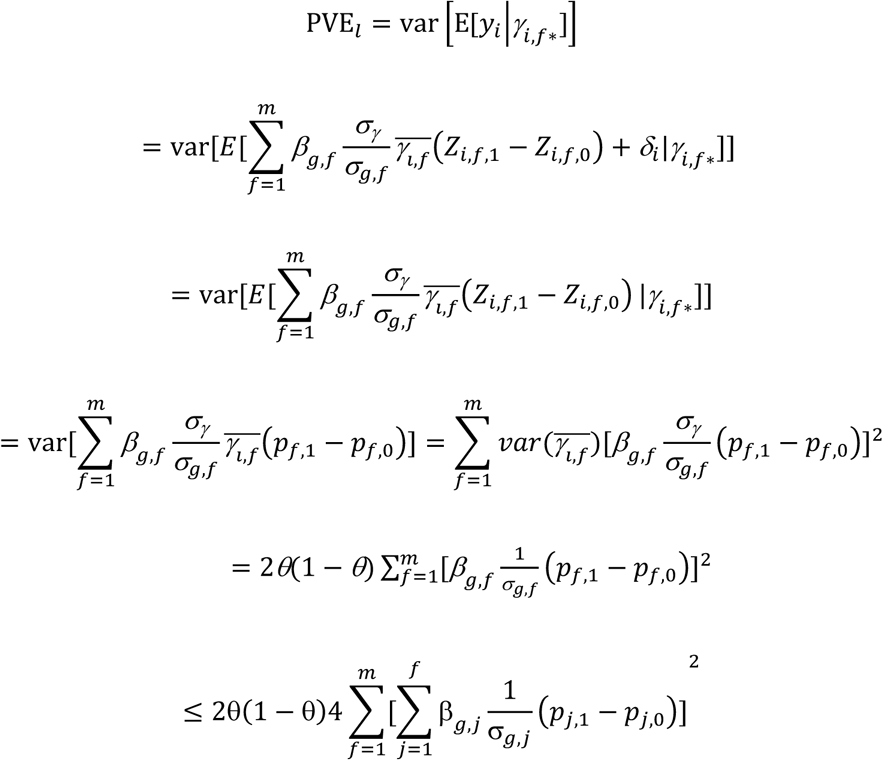

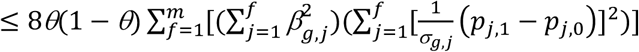

by Cauchy-Schwarz inequality. This implies:

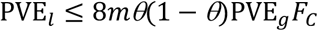

where 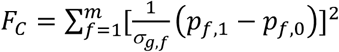 this inequality using simulations (see Additional file 1: **Table S1**). This relates the trait variance explained by local ancestry, the aggregate genetic effect on phenotype, the level of population differentiation of the causal variants, and the degree of polygenicity of the trait. We note that the above derivation assumes that 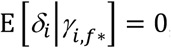, i.e., there are no other causal genetic effects unaccounted for. Equation (2*), as the derivation shows, applies in a more restrictive setting, with the dual assumptions of polygenicity and independence.

Note 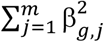 is the aggregate genetic effect and 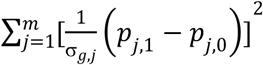 the total extent of population differentiation for the causal eQTLs included in the sum. Since the latter depends on the number of causal eQTLs, it may be useful to consider the mean level of population differentiation in the cis region, 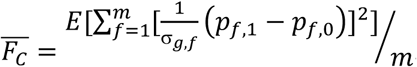.

### Local ancestry, its aggregate effect, and trait heritability estimation

A linear mixed model (LMM) can be used to obtain an aggregate estimate of regulatory (genetic) effect on gene expression. For a given *n*-vector *g* of gene expression levels for *n* individuals, the LMM approach fits the following model:

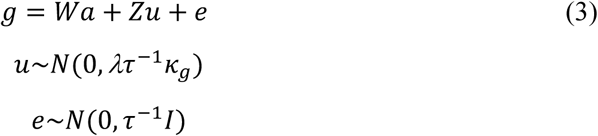

Here *W* is a matrix of covariates (of dimension *n×p*), *a* is the *p*-vector of effects for the covariates (including the intercept term), *Z* is an *n×m* matrix, *u* an *m*-vector of random effects, *e* is the residual vector, *κ*_*g*_ is a genetic similarity matrix, *τ*^-1^ is the variance of residual errors and *λ* is the ratio of two variance components. The approach estimates PVE by genetic variants (PVE_g_) defined as follows:

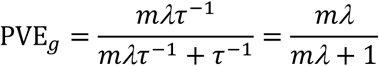

using restricted maximum likelihood (REML). We note that the random genetic effect *Zu* is gene-specific. This Simple-LMM model has been used to characterize infinitesimal genetic architectures in an ancestrally homogeneous population. However, we evaluated the concordance with results from assuming a more general genetic architecture, namely a mixture distribution for the effect sizes, using Bayesian Sparse Linear Mixed Model (BSLMM) [33], which includes the LMM and Bayesian Variable Selection Regression as special instances.

A revised version of the univariate LMM model (3) can be used to estimate the trait variance explained by local ancestry. Here analogously to using the SNP data, the similarity matrix *κ*_*l*_ [24] is constructed from the local ancestry values:

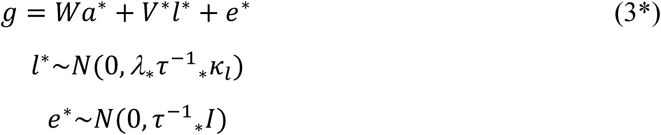

Model (3*) allows estimation of the PVE by local ancestry (PVE_*l*_):

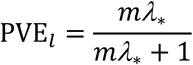

using REML. Alternatively, the effect explained by local ancestry throughout the genome can be modeled to derive from a mixture of a normal distribution and a point mass *δ* at 0:

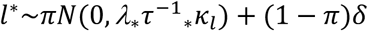

where *π* is the proportion of non-zero effects in the genome. In simulations, we assessed the accuracy of the estimate of PVE_*l*_ from the Gaussian approach (versus a mixture approach) to modeling the effect size explained by local ancestry. Under the same assumptions for equation (2*), model (3*) provides an estimate 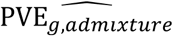 for trait heritability, as also previously noted [24]:

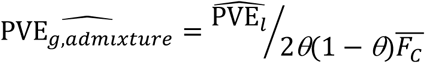

However, in contrast with [24], this estimate is more appropriately viewed as the “expected heritability” in the presence of admixture, departure from which yields additional insights into genetic architecture (i.e., violation of the assumptions of polygenicity and independence) or may indicate presence of stratification. Notably, we also obtain a measure 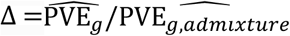 of departure from expectation if Δ is substantially different from one:

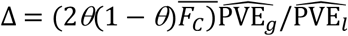

Thus, 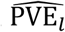 can be used to estimate the expected heritability given admixture, as in the expression for 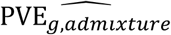, but also to evaluate the potential presence of population stratification due to local ancestry, as in the expression for Δ.

We also implemented a joint model which partitions gene expression into two components, the genetic component (*G*) and local ancestry component (*L*):

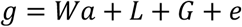

The local ancestry component *L* may be written as a function of the *m* causal variants: *L* = *f*(*×*_1_, *×*_2_, …, *×*_*m*_). A simple estimator is the first principal component derived from the (whole-genome) genotype matrix (i.e., an estimate of global ancestry). Other statistical approaches can be used with varying predictive and computational performance. By explicitly modeling the component due to local ancestry, we may get a more accurate estimate of the overall genetic effects. However, the gain in accuracy depends on the choice for fitting the estimate 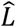. In our approach, which we term Joint Genetics and Local Ancestry (Joint-GaLA), for computational purposes and simplicity, we assume Gaussian distributions for *G* and *L* and restrict the model to the variants in the cis region:

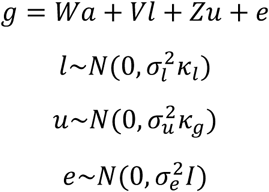

Here *u* and *l* are random effects with corresponding similarity matrices *κ*_*g*_ and *κ*_*l*_ generated from local genetic variation and the corresponding local ancestry, respectively. Using simulations (see next section), we assessed the accuracy of the estimate of 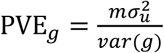 from Joint-GaLA and compared this estimate from that obtained from Simple-LMM (equation (3)). Furthermore, we compared this model with the use of global ancestry to fit 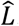.

### Simulation framework for heritability estimation

We conducted extensive simulations, using both real genotype (from the NIGMS AA dataset up to 500 causal variants) and simulated genotype data of admixed samples, in order to (a) validate the analytically derived relationships (inequality (2) and equation (2*)) and confirm the expression for the “expected heritability” in the presence of admixture as well as show departure from the expected value in the presence of local ancestry stratification, (b) evaluate the accuracy of the PVE estimation methods assuming different levels of stratification, and (c) compare the PVE_*l*_ estimate and the *R*^2^ from global ancestry (estimated from simple linear regression).

To simulate genotype data, we assumed 2 ancestral populations, 1000 individuals, 1000 variants in the cis region, *F*_*st*_=0.16 and *F*_*st*_=0.05, and *h*^2^=0.30 (the observed mean in the GTEx skeletal muscle data) and 0.80 for the phenotype. Global ancestry *θ*_*i*_ for the *i*th individual was drawn from a truncated normal distribution *N*(0.7,0.2). Local ancestry at the variant was defined as the sum of two draws from the binomial distribution *Bin*(1, *θ*_*i*_). The ancestral allele frequency was assumed to be distributed as *Unif*(0.05, 0.95) and, with *F*_*st*_, was used to generate the variant allele frequency, which was drawn from the beta distribution with parameters *p*(1 – *F*_*st*_)/*F*_*st*_ and (1 – *p*)(1 – *F*_*st*_)/*F*_*st*_. The genotype for the *i*th individual at the *k*th causal variant was then derived from a random draw from the binomial distribution with expected value defined by the local ancestry for the individual. We assumed 1 local ancestry transition (since local ancestry tract in AA is usually >10 Mb). We varied the number of causal eQTLs (10, 25, 100, 200, 500, 1000) to assess the accuracy of the method as a function of sparsity or polygenicity. (We describe below the simulation framework for the case *m*=1 in the simulations for genetic association [eQTL] mapping.) The effect size of the *k*th causal variant was simulated as *β*_*k*_∼*N*(0, *h*^2^/*m*), with *m* equal to the number of causal variants. As we previously noted, this assignment of effect sizes is a strong assumption (shared with the widely used GCTA or LDSR [21]) about how heritability is distributed among the causal eQTLs and is independent of LD.

We simulated gene expression as follows:

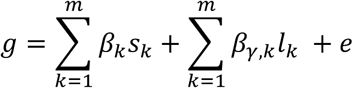

The first summation is the phenotype effect due to genetic variation while the second is due to local ancestry. Because the local ancestry tract typically exceeds the size of the cis region, we assumed a constant value for *l*_*k*_ in the second summation. The single local ancestry effect size 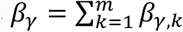 was obtained from the empirical distribution (in NIGMS) at 4 different percentiles (the quartiles for 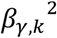 at 0.00138, 0.005444, 0.01755, 0.2877) representing different levels of stratification. The residual *e* was added and assumed to be distributed as 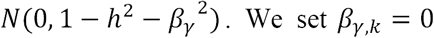. We set *β*_γ,*k*_ = 0 when simulating gene expression without stratification.

We derived estimates from Simple-LMM and Joint-GaLA from 100 independent runs for each set of choices for the parameters. Estimates for PVE_*l*_ were obtained, assuming model (3*), in 100 independent runs to confirm equation (2*).

Departure from the expected heritability 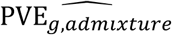 was tested, assuming local ancestry stratification, in simulations with real genotype data (Mann-Whitney U test in 100 independent runs). In this case, we calculated the mean level of *F*_*st*_ (equation (2*)) for the tested causal variants using allele frequency information from the 1000 Genomes CEU and YRI for the ancestral populations.

### Comparison with LD Score Regression for estimate of population stratification and of heritability

LDSR is a widely used approach to estimate confounding due to population stratification and to estimate heritability, using only GWAS summary statistics. We therefore sought to investigate how LDSR performs at these tasks in 100 independent runs for each set of configurations defined as above using *F*_*st*_, *h*^2^, the number of transitions, and *m*. We calculated the LD Score at each variant 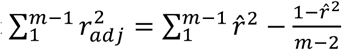 using the LD in the NIGMS genotype data. The use of the actual LD as observed in the dataset simulates the use of a perfectly matched population reference panel. We ran linear regression with simulated gene expression and real genotype data and with global ancestry as a covariate. We applied LDSR to the simulated GWAS datasets to estimate the heritability PVE_*g*_ and the amount of confounding as quantified by the “intercept” (along with the standard error for each). We note that LDSR, by design, does not provide an estimate for PVE_*l*_.

### PVE estimation in real transcriptome data

We estimated the PVE by local genetic variants (PVE_*g*_), defined as within 1 Mb of the gene, for each gene in GTEx skeletal muscle AA samples using REML as implemented in GCTA [34]. We used this tissue in order to maximize the number of AA samples (n=57). We used only common variants (MAF>0.10; n=6,122,246) in this AA subset to increase the estimation accuracy. We calculated the gene-specific genetic relatedness matrix (*κ*_*g*_) using local genetic variants and incorporated 3 PCs, 10 Probabilistic Estimation of Expression Residuals (PEER) variables [27], sex and sequencing platform as fixed effects in the LMM. We used a non-constrained model which allows the PVE estimates to be negative or larger than 1 in order to obtain unbiased estimates but restricted ourselves to genes whose estimates were between 0 and 1 in the downstream analysis. We used the p-value from the likelihood ratio test for the genetic variance component to select genes with nominally significant estimates (nominal p-value<0.05) and Benjamini and Hochberg (BH) corrected [35] FDR<0.10.

We randomly selected 57 samples of European descent out of the 491 GTEx samples in order to compare the PVE_*g*_ between two populations. We selected common variants in this subsect (MAF>0.10; n=4,946,431) and applied the LMM approach described above.

We identified differentially expressed genes between AAs and European Americans (EAs) in skeletal muscle tissue using a *t*-test (BH FDR<0.05).

Similarly, we estimated PVE by local ancestry (PVE_*l*_) at common local variants in the GTEx skeletal muscle AA samples. We used the estimated local ancestry around each gene (within 1 Mb of the gene) to construct the relatedness matrix (*κ*_*l*_). The LMM was fitted to estimate PVE_*l*_ for each gene expression phenotype using the same set of fixed effects as in the PVE_*g*_ analysis.

Using GTEx skeletal muscle AA data, we applied Joint-GaLA (see above). The estimated PVE_*g*_ from Joint-GaLA was then compared with the estimate from the Simple-LMM model.

We investigated the possible reasons for any observed difference in PVE_*g*_ between the populations. We performed LMM association analysis using GEMMA by fitting a model of gene expression with each local SNP and the genetic relatedness matrix constructed from local SNPs. We compared the distribution of allele frequency and of effect size for nominally significant SNPs (p-value<0.05 from the LMM association) between the populations. We also considered the variance in genetic relatedness *A*_*jk*_ generated from the local genetic variants for pairs of distinct individuals:

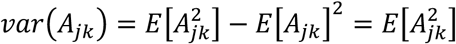

By definition, 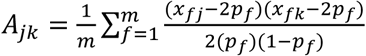, where *x_fj_*is the genotype at variant *f* for individual *j*, *p* _*f*_ is the allele frequency, and *m* is the number of local variants. Now, 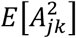 simplifies to the sum of LD correlations over all pairs of variants used in the relatedness matrix [*A*_*jk*_], as has also been previously noted [25]. Thus, the variance in relatedness, *var*(*A*_*jk*_), can be used to evaluate the effect of differential LD pattern near the gene on the population specificity of its genetic regulation.

### Sparsity or polygenicity of gene expression

To systematically characterize the sparsity or polygenicity of gene expression in a recently admixed population, we applied Bayesian Sparse Linear Mixed Model (BSLMM) [36] to generate an estimate of PGE (the proportion of variance explained by the sparse genetic effect) and PVE_*g*, *BSLMM*_ (the sum of the polygenic and sparse effects) for each gene in the GTEx skeletal muscle AA dataset. This analysis would determine genes for which gene expression is influenced by a small number of genetic variants. We calculated the Spearman correlation between PVE_*g*,*BSLMM*_ from BSLMM and PVE_*g*,*lmm*_. We identified genes with highly discordant estimates between the two methods (i.e., PVE_*g*,*A*_ ∉ [PVE_*g*,*B*_ – 2 * *SE*(PVE_*g*,*b*_), PVE_*g*,*B*_ + 2 * *SE*(PVE_*g*,*B*_)], for PVE estimation methods *A* and *B*). We performed simulations (see above) to evaluate the accuracy of the LMM approach as a function of the number of causal variants (i.e., as a function of a sparse or polygenic architecture).

### Use of local ancestry in eQTL mapping

The statistical approach assumes an additive effect of genotype on gene expression and adjusts for the variant-level local ancestry covariate in addition to the sample-level covariates (such as age, sex, or principal components). For each gene-variant pair, we fit the following baseline model:

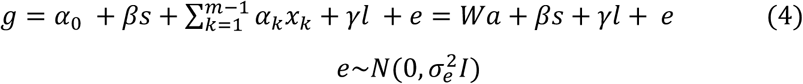

where the *n*-vector *g* is the expression measurement of a gene for the *n* individuals, *s* is the genotype of a marker (typically a SNP proximal to the gene, e.g., within 1 Mb) encoded by 0, 1, and 2 representing the number of alterative alleles with effect size *β* on expression level, *x*_*k*_ is the *k*-th covariate (e.g. age, sex) with effect *x*_*k*_, *x*_0._ is the intercept, *l* is the local ancestry encoded by 0, 1, and 2 according to the number of African ancestry alleles at the tested variant with effect size *γ*, and *e* is the residual assumed to be normally distributed with mean 0 and variance 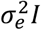. Here *W* is a *nxm* matrix of covariates, including the intercept term, with weight *a*. The baseline model accounts for population structure by adjusting for the local ancestry, while in the usual model, the admixture proportions or, because of computational efficiency, the top PCs of the genotype matrix are incorporated into the model as quantitative covariates (among the *x*_*k*_’s) and locus-specific ancestry is ignored.

Since the genotype *s* and local ancestry *l* at the variant may be correlated, we estimated how much the variances in the effect sizes, *var*(*γ*) and *var*(*β*), may be increased because of multicollinearity. We fit the ordinary least square regression *l*∼*s*, estimated the *R*^2^, and calculated the variance inflation factor *VIF*(*γ*) = 1/(1 – *R*^2^).

We implemented this model, building on a widely-used eQTL mapping method, Matrix eQTL [10]. Matrix eQTL speeds up the eQTL mapping process by performing billions of association tests via matrix operations. The Matrix eQTL algorithm first regresses out the covariates (age, sex, PEER variables, etc.) from each gene expression trait and each genotype and then standardizes residuals to obtain 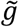 and 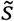, respectively. Then it calculates the correlation 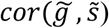 of each residual pair 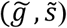 through matrix multiplication and transforms the correlation to a t-statistic 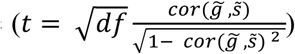.

However, incorporation of local ancestry, which varies by variant, cannot be done in the same manner as the subject-level covariates (e.g. age, sex). Our developed algorithm first regresses out the covariates from gene expression, genotype, and local ancestry to obtain standardized residuals 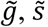, and 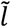. It then regresses out 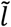 from 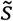 to obtain 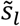 and proceeds to calculate 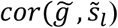 and 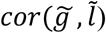 again via matrix operations for efficient processing. We note that equation (4) is equivalent to the following expression after regressing out the covariates:

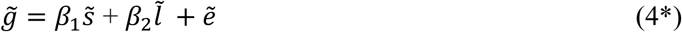

The test for nonzero effect on residual gene expression (*β*_-_ ≠ 0) can be done using an F test (V1 = 1, V2 = N - 2) for the partial correlation coefficient. Equivalently, a t-statistic can be calculated using the following expression:

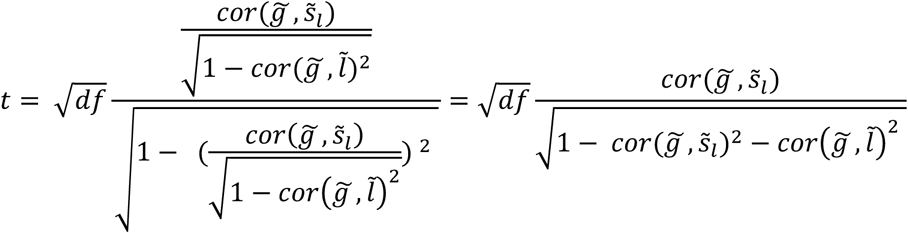

### Type I error simulations for eQTL mapping

In the type I error simulations for eQTL mapping, we considered two scenarios: population stratification due to global ancestry and population stratification due to local ancestry. We utilized real genotype data from the NIGMS AA samples and simulated gene expression levels with different sources of confounding. Because of the difference in variance explained by the first PC and by local ancestry, we utilized the empirical distribution of effect size for each with the scaled expression value (*N*(0,1)) in the NIGMS dataset. We extracted the effect sizes at 4 different percentiles from the empirical effect size distribution for local ancestry and PC, separately analyzed, and used those in the simulations.

We randomly selected 100 genes out of 4595 genes and simulated the gene expression *g*:

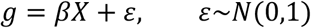

where *β* is the effect size at each percentile and *X* is the first PC or the average local ancestry around each gene. Then we performed cis-eQTL mapping for these genes with no adjustment, global ancestry adjustment (adjustment for 3 PCs), or local ancestry adjustment. We used a range of p-values from 1×10^-6^ to 1 to calculate the false positive rate. We repeated the simulation 1000 times and averaged the false positive rate.

### Type II error simulations for eQTL mapping

In order to test the effect of different population structure adjustment methods on the type II error rate, we first randomly chose 1000 SNPs from the NIGMS genotype data and simulated 500 gene expression variables with standard normal distribution. We tested two scenarios. In the first scenario, the gene expression was only associated with the genotype. We randomly selected 50 SNPs to be true eQTLs whose effect sizes of genotype are 0.9. This choice for the eQTL effect size was motivated by the median of the absolute value of the estimated effect sizes for the significant SNP associations (BH adjusted p<0.05) with scaled gene expression (*g* ∼ *N*(0,1)) in the NIGMS data. In the second scenario, both genotype and local ancestry contributed to the gene expression. We randomly selected 50 SNPs, and effect size of the genotype is 0.9 and that of local ancestry is 0.8. Again, the effect size of local ancestry was chosen from the significant local ancestry associations (BH adjusted p<0.05) with gene expression from fitting a regression *g* ∼ *SNP* + *LA*, where *g* is also the scaled gene expression, in the actual NIGMS data. We performed the eQTL estimation using no adjustment, global ancestry adjustment, or local ancestry adjustment. We used a range of p-values from the minimum to the maximum p-value in each simulation to identify the number of false positives and true positives and calculated the area under the curve (AUC) of the Receiver Operating Characteristic (ROC) curve to summarize the performance of each approach [37]. We repeated the simulations 100 times and plotted the average ROC. We compared the AUC of global ancestry adjustment and local ancestry adjustment for a false positive rate in the range 0-0.2 using a paired two-sided *t*-test.

### Cis-eQTL mapping in NIGMS and GTEx

To identify eQTLs in the NIGMS data, we tested associations between each gene and SNPs within 1Mb upstream of the gene start site and 1 Mb downstream of the gene end site using the local ancestry adjustment approach Joint-GaLA- QTLM. We compared the association results from the adjustment for 1, 2, or 3 PCs and gender and those from the adjustment for local ancestry and gender.

We utilized a hierarchical correction method to identify eQTLs. This method was demonstrated to produce a lower false discovery rate (FDR) and greater true positive rate than the method that applies correction over all association tests [38]. We first used the Benjamini and Yekutieli (BY) procedure [39] to adjust p-values for all association tests by each gene. We then pooled the minimum BY-adjusted p-value of every tested gene to obtain its best associations. We corrected the pooled minimum p-values by the BH correction method [35]. We selected significant eGenes with the threshold of 0.10 for the BY-BH adjusted p-values and used the corresponding minimum BY-adjusted p-value as the threshold to select significant SNPs for these eGenes.

We utilized the eQTL mapping results in the GTEx (v7) LCLs with 117 multiethnic samples as a replication panel. We calculated the replication rate for eQTLs unique to the local ancestry adjustment approach and to the global ancestry adjustment approach.

We then applied Joint-GaLA-QTLM (equation (4)) to the GTEx whole blood and LCL datasets. We excluded samples with East Asian ancestry and used 356 and 114 samples in these two datasets respectively for the cis-eQTL mapping. For the global ancestry adjustment method, we used sex, sequencing platform, 3 PCs, and PEER variables (35 for whole blood dataset, 11 for LCL dataset, consistent with the latest GTEx analysis for the optimal number of PEER factors to avoid overfitting [2]). For the local ancestry adjustment approach, we replaced the 3 PCs with local ancestry. We applied the hierarchical correction method described above and used the threshold of 0.05 for the BY-BH adjusted p-values to select significant eQTLs. We also report the number of eQTLs and eGenes identified at the less strigent threshold (BY-BH p-value<0.1).

We estimated the empirical distribution of the effect size of local ancestry on expression for each gene in both NIGMS and GTEx whole blood datasets while adjusting for the same covariates as in the eQTL mapping.

### Software implementation

We implemented the local ancestry-based eQTL mapping in an R package LAMatrix (https://github.com/yizhenzhong/Local_ancestry). The package incorporates a variant-level covariate (local ancestry) using an algorithm that leverages matrix multiplication to improve computational efficiency.

## Declarations

## Supporting information

## Acknowledgements

We would like to thank Yuan Li and Yinan Zheng for advice on the software development. We thank Tanima De, Zhou Zhang, Yiben Yang and Jun Xiong for helpful discussion. We also thank Alicia Martin for the local ancestry estimation pipeline. E.R.G. benefited immensely from a Fellowship at Clare Hall, University of Cambridge while holding a visiting post in MRC Epidemiology Unit and MRC Biostatistics Unit, Cambridge, UK. We would like to thank the Genotype-Tissue Expression (GTEx) Project, an initiative supported by the Common Fund of the Office of the Director of the National Institutes of Health, and by NCI, NHGRI, NHLBI, NIDA, NIMH, and NINDS, for making the data available to the scientific community.

## Availability of data and materials

The GTEx and NIGMS genotype dataset analyzed during the current study are available in dbGaP under the accession number phs000424.v7.p2 [2] and phs000211.v1.p1[30], respectively. The GTEx expression data are available in GTEx portal (https://www.gtexportal.org/home/) [2]. The NIGMS expression data are included in GEO under the accession number GSE10824 [30].

## Author contributions

YZ performed the analysis, developed implementation, and wrote the software. YZ, ERG, and MP wrote the paper. ERG and MP designed and supervised the analysis, methodology, and data collection for the study. All authors contributed to the interpretation of data. All authors read and approved the final manuscript

## Ethics approval and consent to participate

Not applicable

### Consent for publication

Not applicable

---

Additional file 1: AdditionalFile1.pdf contains Supplementary figures S1-6 and supplementary tables S1-3.

Additional file 2: AdditionalFile2.xls contains PVE estimation in EAs and AAs

Additional file 3: AdditionalFile3.xls contains PVE estimation from LMM model and BLSMM model in AAs

