## Supplementary material for "On Using Local Ancestry to Characterize the Genetic Architecture of Human Phenotypes: Genetic Regulation of Gene Expression in Multiethnic or Admixed Populations as a Model"

**Figure S1. Validation of NIGMS eQTLs for *AMFR* and *PLA2G4C***

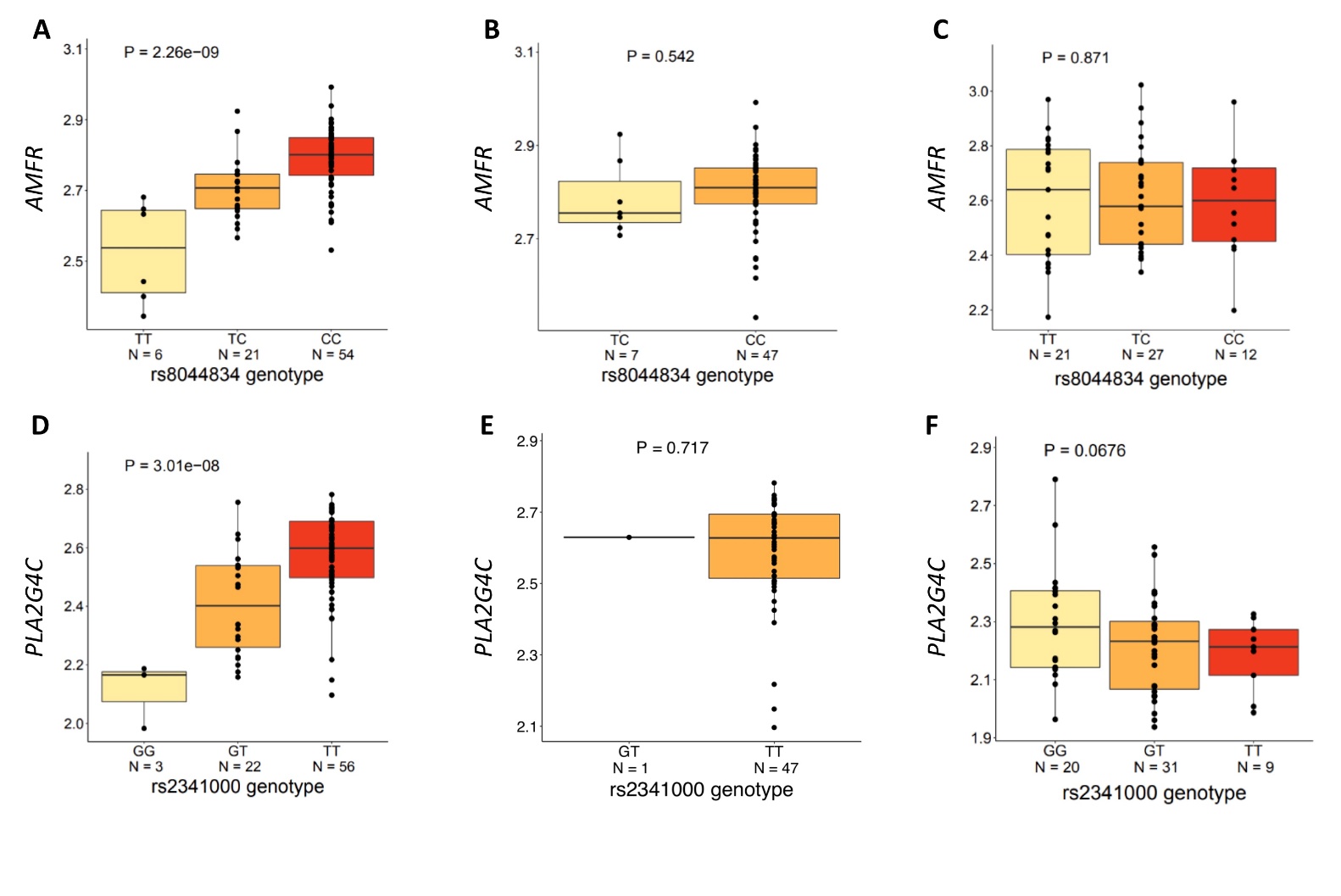

Two eQTLs were found to have highly significant associations by the global adjustment (GA) method, but were non-significant with the local ancestry (LA) adjustment method (purple dots in **Figure 1A**). We investigated whether these eQTLs were driven by association with LA, which may have inflated their p-values with the global adjustment method. Our additional analysis strongly suggests that these were false positive findings.

**A, D.**  Boxplot of genotype to gene expression in NIGMS AA dataset, showing the SNPs as significantly associated with gene expression using the global adjustment approach.

**B, E.** Boxplot of genotype to gene expression in subsamples of two African ancestry alleles in NIGMS AA dataset, showing no support for the associations in an African background.

**C, F.**  Boxplot of genotype to gene expression in HapMap CEU dataset, showing no support for the associations in European background.

**Figure S2. Comparison of results from local ancestry adjustment with results from alternative methods**

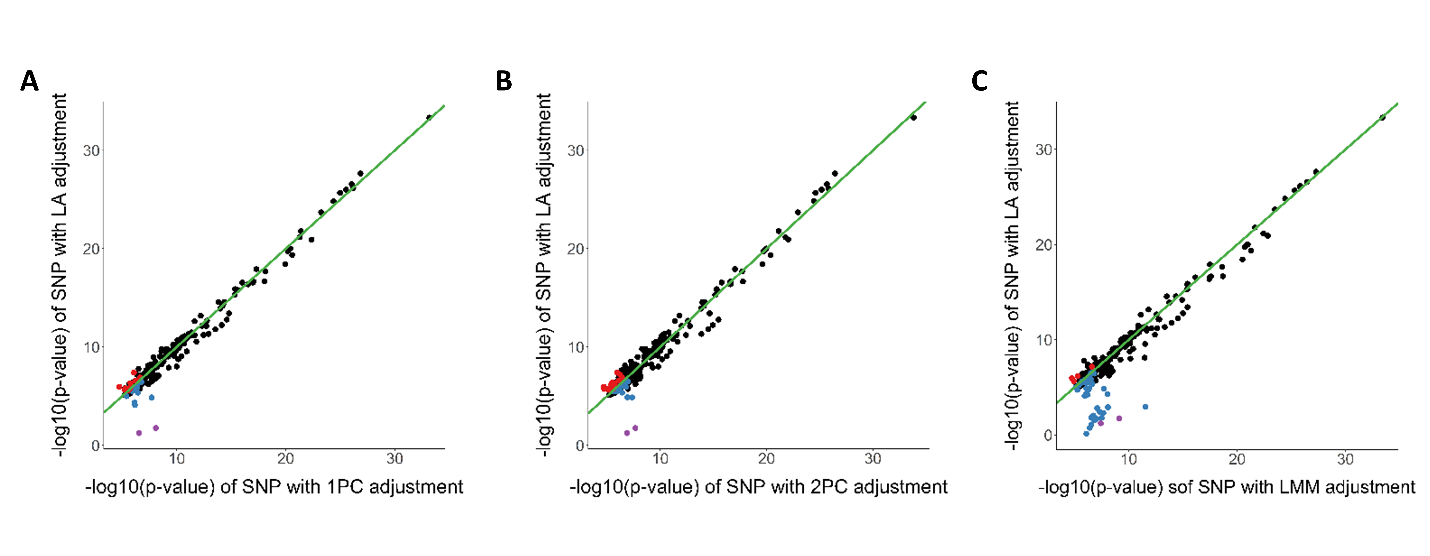

Comparison of results from local ancestry adjustment with results from 1PC adjustment (**A**), 2PCs adjustment (**B**) and LMM method in NIGMS dataset. Results from 1PC and 2PCs adjustment are similar to the results from 3PCs adjustment (**Figure 3A**). LMM mixed model has larger power by identifying more unique eQTLs but fail to remove spurious eQTLs (purple dots).

**Figure S3.** $\mathrm{PVE}_{g}$ **simulations with simulated genotype**

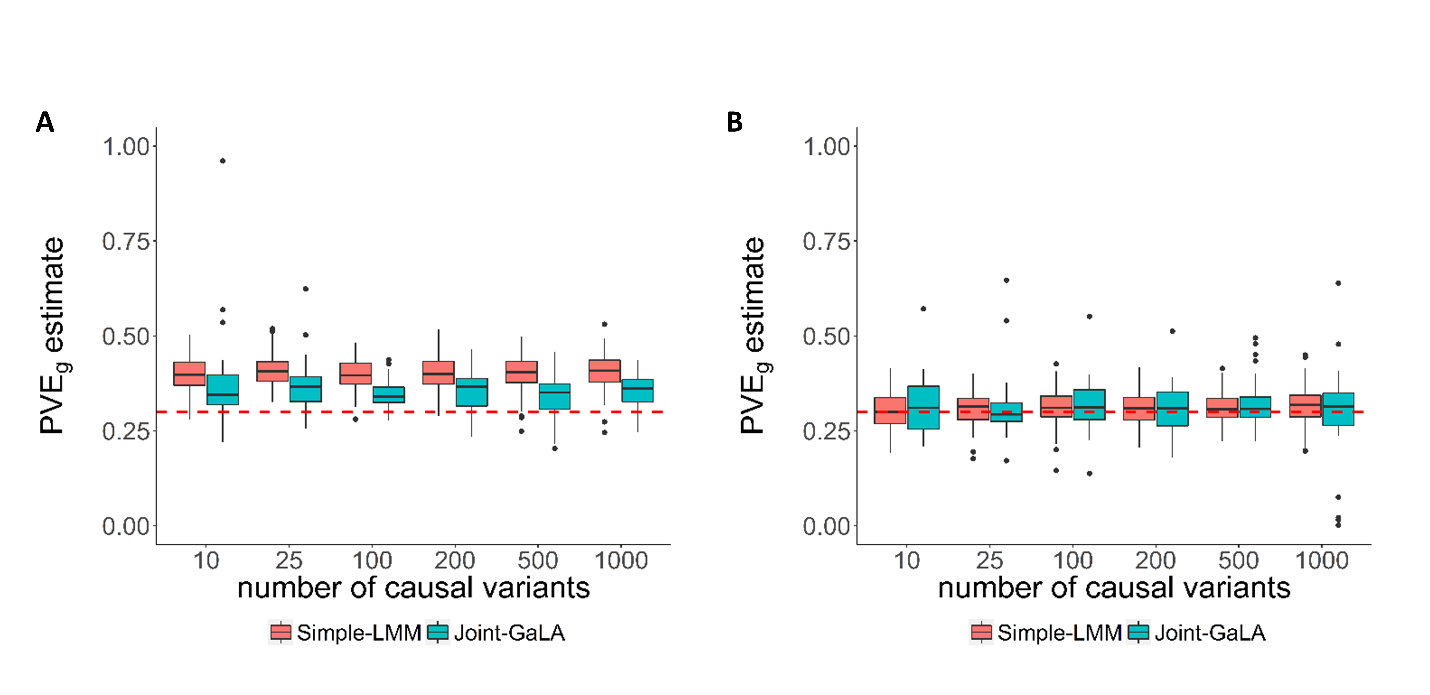

$\hat{\mathrm{PVE}_{g}}$ estimates derived from either simple-LMM or Joint-GaLA even after controlling for global ancestry could suffer from some bias (**A,** $\mathrm{PVE}_{l}=0.2$), but the bias was diminished with a reduction in local ancestry effect on phenotype (**B,** $\mathrm{PVE}_{l}=0.01755$).

**Figure S4. Comparison of** $\mathbf{PVE}_{\boldsymbol{g}}$**estimation in AAs and EAs, related to Figure 6B**

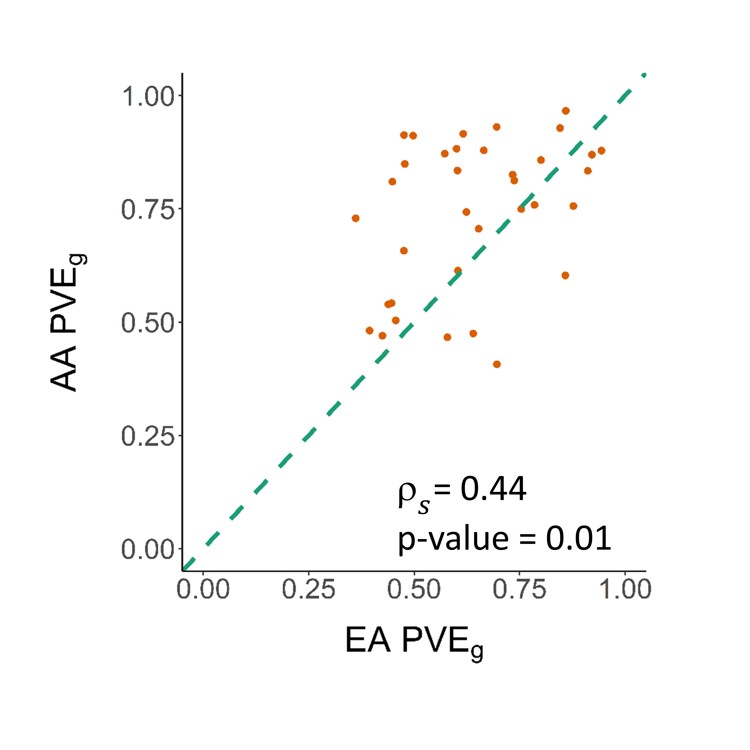

When using FDR<0.1 to select genes with reliable estimates, we continued to see a significant correlation in estimates between EAs and AAs.

**Figure S5.** $\mathbf{PVE}_{\boldsymbol{g}}$**estimation with different methods in AAs**

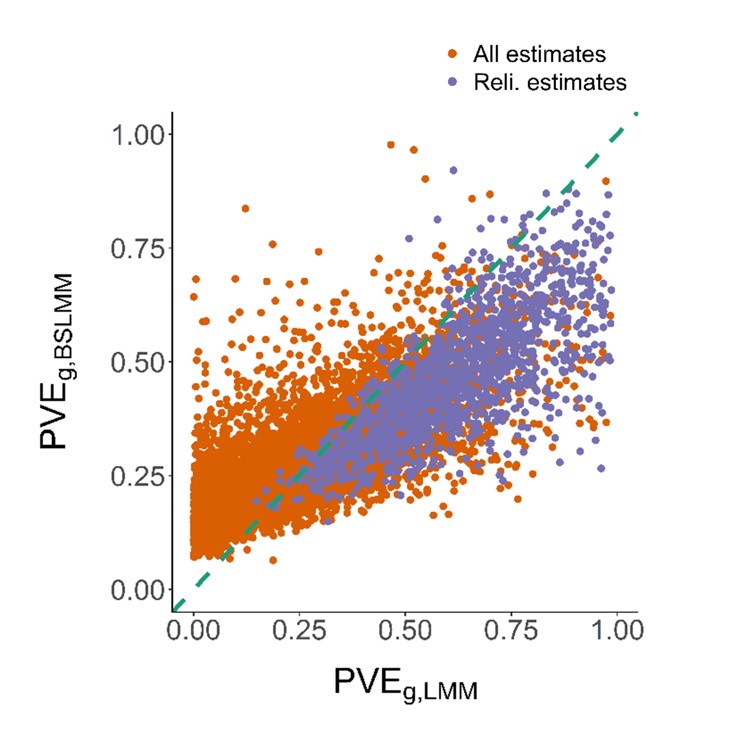

Comparison of $\mathrm{PVE}_{g}$ estimation from LMM and BSLMM model, showing significant correlation (Spearman’s ρ=0.82, p-value<2.2×10^-16^). Reliable estimates were defined as LMM nominal p-value<0.05.

**Figure S6. Correlation of first PC with the average local ancestry across the genome**

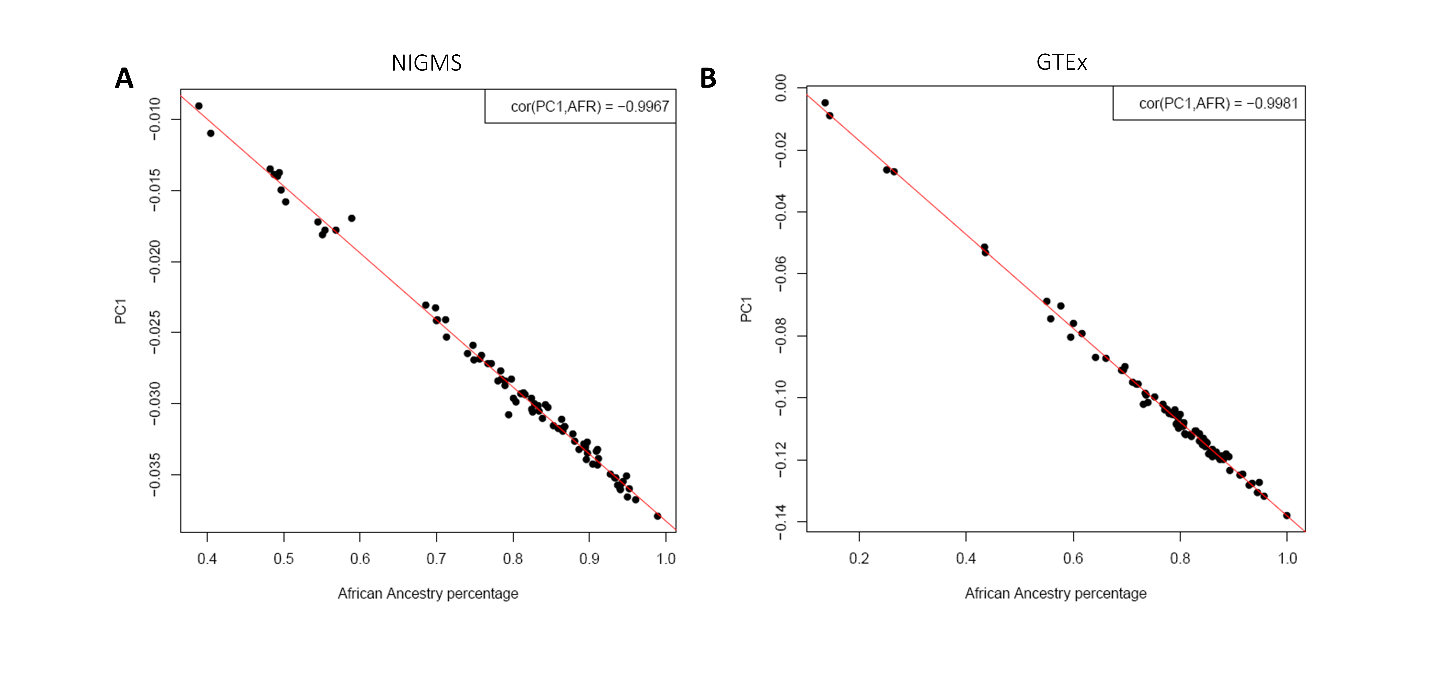

Comparison of the first PC and average African local ancestry (AFR) across the genome in NIGMS (A) and GTEx (B), showing nearly perfect correlation. With decreased EA ancestry (PC1), we see a significant increase in AFR.

**Table S1. Identified eQTLs in GTEx dataset by different methods**

|  | **BY-BH p-value<0.05** | | **BY-BH p-value<0.1** | |
| --- | --- | --- | --- | --- |
|  | GA eQTL(eGenes) | LA eQTL(eGenes) | GA eQTL(eGenes) | LA eQTL(eGenes) |
| $whole blood$ | 840,884(4,963) | 842,476(4,952) | 948,521(5,288) | 950,920(5,292) |
| $\mathrm{LCL}$ | 217,230(1,728) | 225,954(1,777) | 250,150(1,937) | 257,306(1,958) |

**Table S2. PVE analysis**

| PVE analysis | Pop |  | Sample size | Mean  $\mathrm{PVE}_{g}(var)$ | num/percentage FDR<0.1 | num/percentage pvalue<0.05 | num of genes with estimates |
| --- | --- | --- | --- | --- | --- | --- | --- |
| $\mathrm{PVE}_{g}$ | AA |  | 57 | 0.299 (0.051) | 78/0.88% | 1,366/15.47% | 8,832 |
| $\mathrm{PVE}_{g}$ | EA |  | 57 | 0.251 (0.039) | 479/5.52% | 1,703/19.64% | 8,670 |
| $\mathrm{PVE}_{l}$ | AA |  | 57 | 0.298 (0.078) | 40/2.30% | 379/21.78% | 1,740 |

The summary of PVE analysis in GTEx skeletal muscle dataset.

**Table S3. PVE simulation with simulated genotype**

| h^2^ | Number of causal variants | $\mathrm{PVE}_{g}$ Mean±sd | $\mathrm{PVE}_{l}$ Mean±sd | R-squared (GA) |
| --- | --- | --- | --- | --- |
| 0.8 | 10 | 0.796±0.024 | 0.078±0.089 | 1.272e-3 |
|  | 25 | 0.803±0.024 | 0.059±0.039 | 9.727e-4 |
|  | 100 | 0.802±0.023 | 0.061±0.036 | 1.280e-3 |
|  | 200 | 0.799±0.022 | 0.059±0.036 | 7.881e-4 |
|  | 500 | 0.800±0.025 | 0.066±0.050 | 9.026e-4 |
|  | 1000 | 0.799±0.020 | 0.068±0.044 | 1.003e-3 |
| 0.3 | 10 | 0.302±0.044 | 0.037±0.038 | 1.041e-3 |
|  | 25 | 0.304±0.053 | 0.025±0.024 | 8.985e-4 |
|  | 100 | 0.302±0.043 | 0.035±0.084 | 1.016e-3 |
|  | 200 | 0.296±0.049 | 0.023±0.017 | 1.203e-3 |
|  | 500 | 0.301±0.051 | 0.023±0.020 | 9.077e-4 |
|  | 1000 | 0.306±0.054 | 0.042±0.069 | 9.063e-4 |

Using simulations, we confirmed the inequality 2: $\mathrm{PVE}_{l}\leq8m\theta\left( 1-\theta\right)\mathrm{PVE}_{g}F_{C}$.

We also confirmed the equality 2*, thus validating the expected heritability in the presence of admixture: $\mathrm{PVE}_{l}=2\theta(1-\theta)\mathrm{PVE}_{g}\bar{F_{C}}$. The right-hand side is 0.053 ($h^{2}$=0.80) and 0.020 ($h^{2}$=0.30) according to the equation and is close to the estimated$\mathrm{PVE}_{l}.$
